# HCS-3D*X*, a next-generation AI-driven automated 3D-oid high-content screening system

**DOI:** 10.1101/2024.07.15.603536

**Authors:** Akos Diosdi, Timea Toth, Maria Harmati, Istvan Grexa, Bálint Schrettner, Nora Hapek, Ferenc Kovacs, Andras Kriston, Krisztina Buzas, Francesco Pampaloni, Filippo Piccinini, Peter Horvath

**Affiliations:** Synthetic and Systems Biology Unit, HUN-REN Biological Research Centre (HUN-REN BRC), Szeged, Hungary; Single-Cell Technologies Ltd, Szeged, Hungary; Doctoral School of Biology, University of Szeged, Szeged, Hungary; Department of Immunology, University of Szeged, Szeged, Hungary; Buchmann Inst. for Molecular Life Sciences, Goethe-Universität Frankfurt am Main, Frankfurt am Main, Germany; IRCCS Istituto Romagnolo per lo Studio dei Tumori (IRST) “Dino Amadori”, Meldola (FC), Italy; Department of Medical and Surgical Sciences (DIMEC), University of Bologna, Bologna, Italy; Institute of AI for Health, Helmholtz Zentrum München, Neuherberg, Germany; Institute for Molecular Medicine Finland, University of Helsinki, Helsinki, Finland

**Keywords:** 3D cell cultures, Multicellular 3D tumour models, Micromanipulation, High-content screening, Artificial intelligence, Automated light-sheet microscopy, 3D single-cell analysis

## Abstract

Self-organized three-dimensional (3D) cell cultures, collectively called 3D-oids, include spheroids, organoids and other co-culture models. Systematic evaluation of these models forms a critical new generation of high-content screening (HCS) systems for patient-specific drug analysis and cancer research. However, the standardisation of working with 3D-oids remains challenging and lacks convincing implementation. This study develops and tests HCS-3D*X*, a next-generation system revolutionising HCS analysis in 3D imaging and image evaluation. HCS-3D*X* is based on three main components: an automated Artificial Intelligence (AI)-driven micromanipulator for 3D-oid selection, an HCS foil multiwell plate for optimised imaging, and image-based AI software for single-cell data analysis. We validated HCS-3D*X* directly on 3D tumour models, including tumour-stroma co-cultures. Our data demonstrate that HCS-3D*X* achieves a resolution that overcomes the limitations of current systems and reliably and effectively performs 3D HCS at the single-cell level. Its application will enhance the accuracy and efficiency of drug screening processes, support personalised medicine approaches, and facilitate more detailed investigations into cellular behaviour within 3D structures.

## Introduction

For decades, evaluation of drug effects has relied on two-dimensional (2D) cell cultures as model systems. However, 2D cell cultures cannot accurately capture the complex physiological characteristics of tissues and tumour microenvironments^1^. Three-dimensional (3D) cell cultures are gaining more attention as a relevant model system for drug testing^2–4^. In particular, the so-called “3D-oids” (3D models including spheroids, organoids, tumouroids, and assembloids) have been broadly tested and proven to mimic *in vivo* conditions. Indeed, 3D-oids can maintain tissue structure which makes them highly relevant for numerous biological research areas, including drug discovery, regenerative medicine, tumour biology, and immunotherapy^5–7^.

Since 2010, the continuously increasing attention on 3D cell cultures has resulted in almost 40,000 articles (based on the search terms “spheroid, organoid, tumoroid, and assembloid”) and more than 168,000 publications discussing high-content systems. However, only 1% of these publications present high-content screening (HCS) analyses based on imaging systems, and only 76 contributions refer to HCS imaging of 3D-oids (**Supplementary Table 1**). These small numbers show that there are challenges in developing 3D HCS platforms, including 3D-oid generation, handling, imaging, and analysis.

At the level of 3D-oid generation, there are concerns about: (*a*) morphological variability^8,9^, (*b*) penetration properties of compounds, including specific stains^10^, (*c*) inner distribution and biological characteristics of the cells^11,12^. While seminal work has been conducted on standardising generation protocols^13^, the nature of 3D models means that high variability is present when screening large numbers of 3D-oids^14^. To handle 3D-oids while ensuring experimental reproducibility and standardisation, the use of Artificial Intelligence (AI)-driven systems was proposed in order to manipulate and select similar 3D-oid aggregates^15–17^. For instance, AI-driven micromanipulators combining morphological pre-selection with automated pipetting systems reduce time and ensure reliability when transferring spheroids to the imaging plates^18^.

Complex drug screening analyses require single-cell phenotyping, necessitating imaging at the highest resolution^19^. Light-sheet fluorescence microscopy (LSFM) is able to visualise large samples at the cellular level with high imaging penetration, minimal phototoxicity and photobleaching^20,21^. The available LSFM HCS systems for 3D-oids have different drawbacks (*e.g*., different sample preparation, light penetration, imaging time) (**Supplementary Table 2**). In addition, the amount of generated data is typically vast and heterogeneous, making data analysis a time-consuming and computationally demanding procedure requiring automation^22^. Several pipelines and tools have been introduced for the analysis of 3D data, but no standard has been defined for quantitative tasks, including segmentation, classification, and feature extraction^23–25^.

In this work, we present HCS-3D*X*, a customisable HCS system for 3D imaging and analysis of 3D-oids at a single-cell level (**Fig. 1a**). HCS-3D*X* includes (*I*) selection and transfer of morphologically homogeneous 3D spheroids using a custom-developed tool, called *SpheroidPicker* (**Fig. 1b**); (*II*) single-cell LSFM imaging using a custom Fluorinated Ethylene Propylene (FEP) foil multiwell plate (**Fig. 1c, f-k**); (*III*) an AI-based custom 3D data analysis workflow developed in *Biology Image Analysis Software* (*BIAS*, Single-Cell Technologies Ltd., Szeged, Hungary)^26^ (**Fig. 1d**). We present a multitude of experiments to show that HCS-3D*X* can be reliably used for single-cell 3D HCS. First, morphological analysis of tumour spheroids under different conditions was executed to define the best setup for 3D-oid selection using the *SpheroidPicker*, an AI-guided 3D cell culture delivery system^18^. Then, the accuracy of 2D features at multiple levels was validated (**Fig. 1e**). Before extending the feature analysis to 3D cultures, we specifically quantified screening performance and image quality (see below) while using the designed HCS foil multiwell plate. Finally, the effectiveness of the HCS-3D*X* system was proven on monoculture and co-culture tumour models via quantitative evaluation of tissue composition at single-cell resolution (**Fig. 1l-n**).

**Fig. 1:**
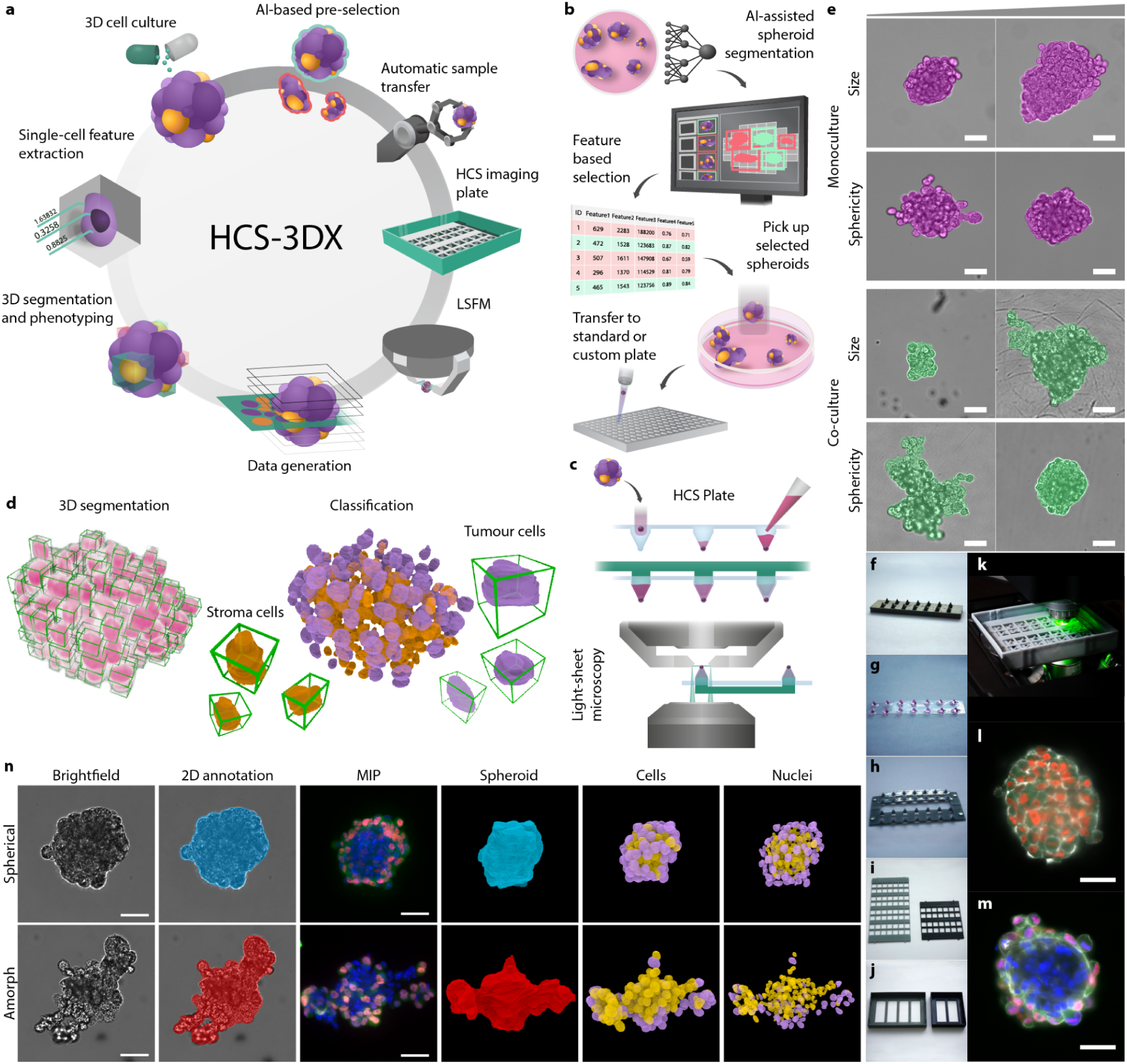
HCS-3D*X*, an AI-driven HCS system for 3D imaging and analysis of 3D-oids at a single-cell level. (**a**) HCS-3D*X* workflow includes 8 essential steps: (*I*) 3D cell culture generation, (*II*) AI-driven 2D object detection, feature analysis and selection, (*III*) automated sample transfer from the source culture vessel, (*IV*) transfer to a custom-developed HCS imaging plate, (*V*) LSFM screening, (*VI*) generation of massive 3D image datasets, (*VII*) image analysis with *BIAS* (*VIII*) feature extraction and single-cell classification. (**b**) Step *II-IV* - working principle of the *SpheroidPicker*, an AI-guided, 3D cell culture delivery system using a light microscope for 2D morphology-based spheroid screening and analysis, enabling semi- or fully-automated transfer of selected spheroids with predefined morphology features. (**c**) Step *IV* - selected spheroids are transferred to the HCS foil multiwell plate compatible with LSFM to obtain single-cell resolution in 3D. (**d**) Step VII - a representative image of a nuclei-labelled spheroid analysed and visualised by BIAS, with single cells indicated by green bounding boxes. (**e**) Representative images of the 2D annotated dataset including monoculture (purple) and co-culture (green) spheroids with the smallest and largest volumes and least and greatest sphericity values. (**f**) 3D printed heat-resistant mould element that was used to form the FEP foil. (**g**) Vacuum-formed transparent FEP foil where each cuvette is suitable for one sample. For visualisation, each cuvette was filled with cell culture medium. (**h**) The insert element that fits into the cuvettes of the foil. The insert element only secures the position of the samples at the bottom of the cuvettes. (**i**) The grid element that fits into the base prevents the movement of the FEP foil after filling up with the detection solution. Two versions are available. (**j**) The base of the plate is an additional component that provides volume for the detection solution and space inside the plate for the water immersion objectives. Two versions are available. (**k**) Assembled HCS plate in use. (**l**) Representative image of a monoculture spheroid composed of HeLa Kyoto cells (green - EGFP-alpha-tubulin; red - H2B-mCherry; grey - actin). The scale bar represents 50 µm. (**m**) Representative image of a co-culture spheroid comprising HeLa Kyoto and MRC-5 cell lines (blue - DAPI, green - EGFP-alpha-tubulin; red - H2B-mCherry; grey - actin). The scale bar represents 50 µm. (**n**) Representative co-culture spheroids showing a spherical and an irregular example from the dataset. Images from left to right show the brightfield images with the corresponding annotation, the maximum intensity projection (MIP) images acquired with the light-sheet microscope, whole spheroid segmentation, and cytoplasm and nucleus segmentation combined with machine learning-based classification. The MRC-5 cells are illustrated with orange, while the HeLa Kyoto cells are illustrated with purple colour. The scale bar represents 50 µm.

## Results

### Concept of HCS-3D*X*

The developed HCS-3D*X* platform overcomes many limitations of the current analysis of 3D-oids. HCS-3D*X* comprises all the steps required to evaluate 3D cell cultures at a single-cell level by enabling reliable, fast and automatic high-content imaging of multiple 3D-oids with a high penetration depth (**Supplementary video 1**). Consequently, the proposed concept is advantageous for a wide range of research purposes, including industrial drug screening, personalised medicine, or basic research.

### 2D features variability facilitates definition of 3D-oid pre-selection parameters

In the first experiment, a comparative study was executed for the pre-selection of spheroids generated under the same conditions. To verify the ideal screening parameters in 2D, brightfield images obtained by various objectives were compared by measuring radiomic features of co-culture spheroids. The same spheroids were imaged at different magnifications (**Supplementary Table 3**). Specifically, 50 spheroids were imaged with 2.5x, 5x, 10x, and 20x objectives resulting in 200 images (**Supplementary Fig. 1a**). The images were manually annotated and 2D morphological features were extracted using *BIAS* and *ReViSP* (a specific tool for estimating the volume of spheroids using a single 2D brightfield image^27^). On average, for all the extracted features (*i.e*., *Diameter*, *Perimeter*, *Area*, *Volume 2D*, *Circularity*, *Sphericity*, and *Convexity*), the relative differences between different magnifications (considering 20x as a reference) were less than 5%, except for *Volume* that reached almost 6.5% (**Fig. 2a** and **Supplementary Table 4**). *Perimeter*, *Sphericity*, *Circularity*, and *Convexity* showed significant differences between the 2.5x and the 5x/10x objectives since the lower image resolution resulted in a less accurate representation (**Supplementary Fig. 1b,c**). The 2.5x objective resulted in the fastest imaging including finding and focusing, however, it showed the least accurate feature extraction. Both 5x and 10x objectives were ideal for imaging spheroids since they increased the imaging speed by ∼45% or ∼20% while providing relatively accurate values. The 20x objective was the most accurate, but it required more time to find and focus on the spheroid. However, to provide the most accurate comparison, the 20x objective was used as a reference for all further experiments.

**Fig. 2:**
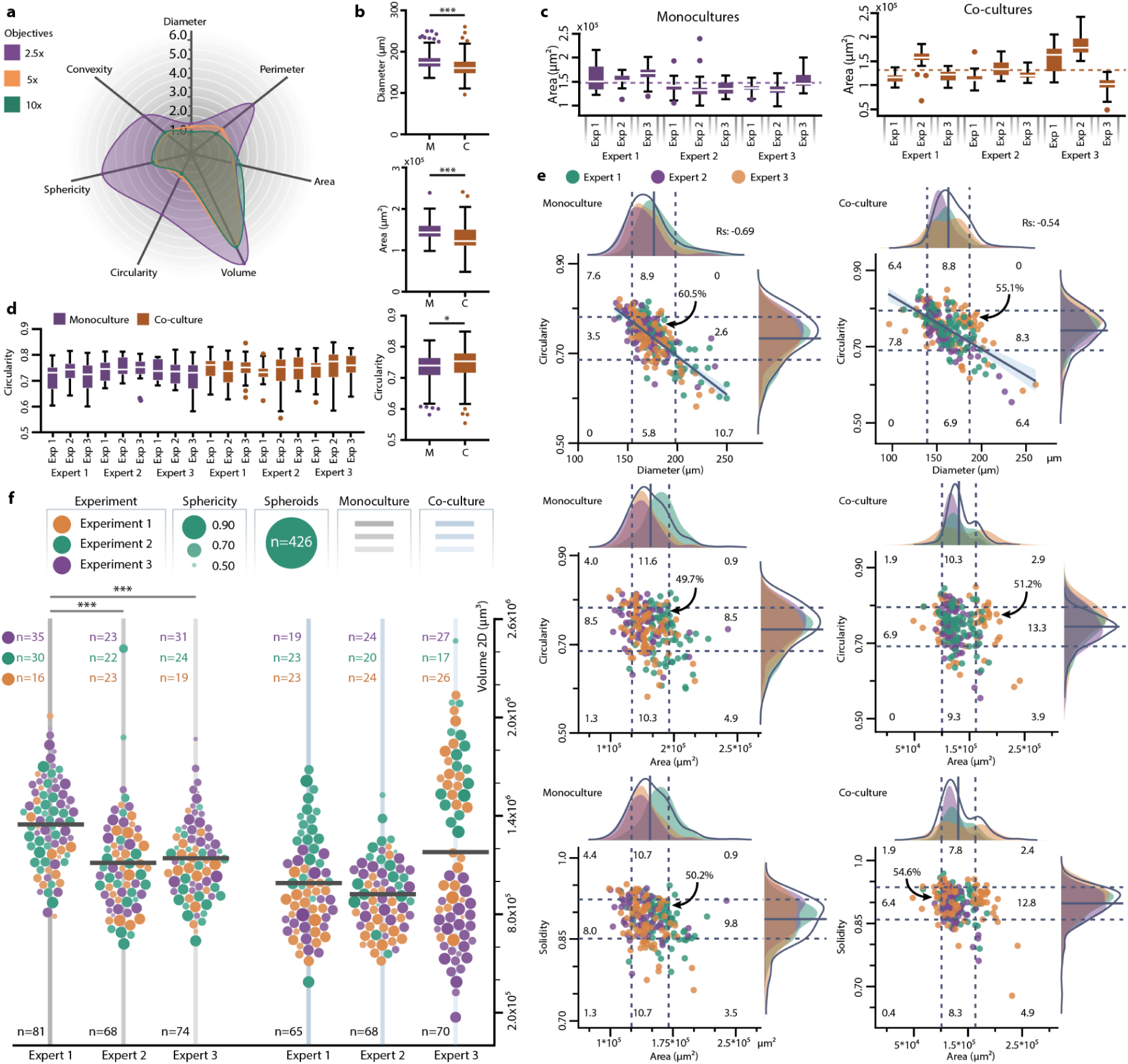
Characterization of monoculture and co-culture datasets using 2D features. (**a**) Spider plot representation of the relative differences of features displayed as a percentage, acquired by annotating the same spheroids, imaged by 4 different objectives with different magnifications. Each objective was compared to the annotations that were executed on the images acquired with the 20x objective. After manual annotation, features such as *Diameter*, *Perimeter*, *Area*, *Volume*, *Circularity*, *Sphericity*, and *Convexity* were displayed (n=50). (**b**) Boxplot visualisation of the monoculture (purple, n=223) and co-culture (orange, n=203) spheroids based on *Diameter* (µm), *Area* (µm^2^), and *Circularity* (−). (**c**) Size comparison of spheroids for each experiment (n≥16). Significant differences are not visualised. Dashed lines show the overall average of monoculture and co-culture experiments. (**d**) Shape comparison of spheroids for each experiment (n≥16). (**e**) Distribution of monoculture (n=223) and co-culture (n=203) spheroids based on the *Diameter*, *Area*, *Solidity*, and *Circularity* features. Each circle represents a spheroid, and each colour corresponds to different experts (green - *Expert 1*, purple - *Expert 2*, and orange - *Expert 3*). The central lines display the average value for all the samples whilst the dashed lines display the plus and minus one standard deviation. Based on the standard deviation, 9 different regions with a corresponding percentage of the number of spheroids are shown. Spearman correlation was used to evaluate the relationship between the variables. (**f**) Beeswarm plot of the monoculture (grey) and co-culture (light blue) spheroids divided by experts (vertical distribution) and experiments (orange - *Experiment 1*, green - *Experiment 2*, purple - *Experiment 3*), indicating the volume and sphericity of each sample. Overall, 426 spheroids were screened. Each circle represents a spheroid on a volume axis showing the sphericity values by the size and intensity (bigger and more intensive circles represent the most spherical spheroids). The number of spheroids evaluated in each batch is displayed, with different colours representing individual experiments and black indicating the total count. For the statistical analysis, a non-parametric Kruskal-Wallis test was conducted, followed by Dunn’s multiple comparison test. When comparing only two groups, the Kolmogorov-Smirnov test was utilized. *p ≤ 0.05; **p ≤ 0.01; ***p ≤ 0.001. Abbreviations: M - monoculture, C - co-culture, Exp - experiment.

### Analysis of spheroid model variability

In order to measure tumour model heterogeneity, mono- and co-culture spheroids generated by 3 experts with extensive daily experience in 3D cell cultures were compared. The experts repeated the same experiments 3 times (**Fig. 2b-f**). A total of 426 spheroids, 223 mono- and 203 co-cultures were generated. For the monoculture spheroids, each expert generated samples of 100 HeLa Kyoto human cervical cancer cells per well in a 384-well U-bottom cell-repellent plate. In the case of co-cultures, each spheroid contained 40 HeLa Kyoto cells and 160 MRC-5 human fibroblast cells representing the tumour stroma. Each spheroid was manually annotated to extract the 2D features.

Although all the experts used the same equipment within the same environment and followed exactly the same protocol, the inter-operator comparison showed great variability regarding the size and shape of the generated spheroids (**Fig. 2c-f**). The direct comparison of mono- and co-culture spheroids showed significant differences while evaluating *Diameter*, *Circularity*, and *Area* (**Fig. 2b**). By evaluating the 2D features of the monoculture spheroids, *Expert 1* generated significantly bigger spheroids (**Fig. 2c,f**). Although the range of *Area* and *Volume (2D)* is wide, *Circularity* and *Sphericity (2D)* values were independent of spheroid size and there were no significant differences between the experts and batches (**Fig. 2d,f**). We observed increased variability between experts and between batches when generating the co-culture spheroids with two different cell lines (**Fig. 2e**). Compared to monocultures, the co-culture spheroids had twice as many seeding cells and, on average, smaller *Volume 2D*, indicating more compact spheroids. Visualising the individual spheroids based on 2D parameters and their respective standard deviation revealed the distribution and the number representing the most similar samples differed (**Fig. 2e**). The data showed stronger correlation and a greater number of spheroids when arranged according to *Circularity* and *Diameter*: −0.69 and 60.5% for monoculture and −0.54 and 55.1% for co-cultures. Using *Area* instead of *Diameter* resulted in no correlation. Furthermore, spheroids were less similar resulting in a smaller quantity of spheroids for *Circularity* and *Area* and *Solidity* and *Area* (49.7% and 50.2% for monoculture and 51.2% and 54.6% for co-culture). Overall, feature selection for spheroid characterization is crucial in terms of outcomes. In particular, evaluations should include one morphological and one size-related feature.

### Design of an optimized HCS imaging plate for LSFM

An HCS multi-well plate with predefined positions for the samples was designed to allow fast and automatic multiple object screenings using an LSFM (**Fig. 1f-k** and **Supplementary Note 1**). The plate was developed and tested on the Leica TCS SP8 DLS upright LSFM (**Supplementary Fig. 2a-f**). To validate the resulting image quality and measure the screening time, T-47D spheroids were generated with a diameter under 200 µm. Two different groups of 5 spheroids were defined: as recommended by Leica, the first 5 were embedded into agarose in U-shape glass capillaries inserted in Petri dishes; the other 5 spheroids were placed into the HCS plate. Regarding the image quality, the difference between the groups was minimal and not noticeable to the naked eye (**Supplementary Fig. 2a**). Therefore, we measured the quality of the images using the intensity variance metric, which can characterise the general blurriness of an image qualitatively^28^. Considering the average values, no significant differences were observed, suggesting similar image quality (**Supplementary Fig. 2b**). Although the spheroids screened with the HCS plate showed slightly better scores, this phenomenon can be explained by the size and morphology differences of randomly selected spheroids.

Next, we analysed the effectiveness of the HCS plate. To measure the average screening time, 10 spheroids were separated into Petri dishes and embedded into agarose, while the other 10 spheroids were placed into the HCS plate. Comparing the total screening time, the HCS plate performed twice as fast (∼50±7 min) while the screening time in the Petri dishes took approximately 102±9 min (**Supplementary Fig. 2c**). The largest difference was observed during sample preparation: manually plating the spheroids into the HCS plate took 14±2 min, while Petri embedding the spheroids into agarose took approximately 37±5 min. Another significant difference appeared during the calibration process, where replacing and calibrating each spheroid took approximately 26±4 min. This does not apply to the HCS plate since all 10 spheroids were separated and placed into individual cuvettes on the same plate. Other calibration and imaging processes took 30±3 min for both approaches. However, finding samples within the plate is easier, as all positions are predefined and one calibration at the beginning is sufficient. Thus, our HCS plates achieved image quality that matched previous studies^28,29^ while showing greater effectiveness that will optimize screening. Overall, the custom-developed HCS plate is easy to use, reduces imaging times, and ensures the same image quality as the method recommended by the manufacturer. The plate is compatible with fixed and optically cleared spheroids with a diameter of 350 µm (**Supplementary video 2**), live-cell imaging of hydrogel-based multi-cellular models, and creating high quality 3D datasets (**Supplementary Fig. 2d-f** and **Supplementary video 3**).

### 2D and 3D feature comparison

From the spheroids used for the 2D imaging, 110 monoculture and 114 co-culture spheroids were randomly selected, and fluorescence images were acquired by LSFM using the HCS plate. Images were imported to the BIAS software to measure 3D features at the nucleus, cell, and spheroid levels (**Supplementary Note 2 and 3**). First, the correlations were analysed between the 2D and 3D features independently for each dataset (**Fig. 3a**). Among the tested features, *Solidity* showed the highest correlation when compared to other shape features both 2D and 3D. There was no correlation between the size and shape descriptors. In general, the 2D estimation resulted in smaller spheroids, while each shape descriptor reached higher values and showed a significant difference compared to 3D. Monoculture spheroids showed a significant difference between *Volume 2D* and *Volume 3D*, but there was no significant difference for the co-culture spheroids. Percentage differences from the average values of the 2D and 3D features were measured where *Volume* and *Solidity* reached the lowest values (18.6% and 18.8% for the monoculture; 4% and 21.4% for the co-culture) (**Fig. 3c**). Although the co-culture spheroids showed a particular 4% difference for the direct comparison, the whole dataset showed a 19.6% difference **(Supplementary Fig. 3b**).

**Fig. 3:**
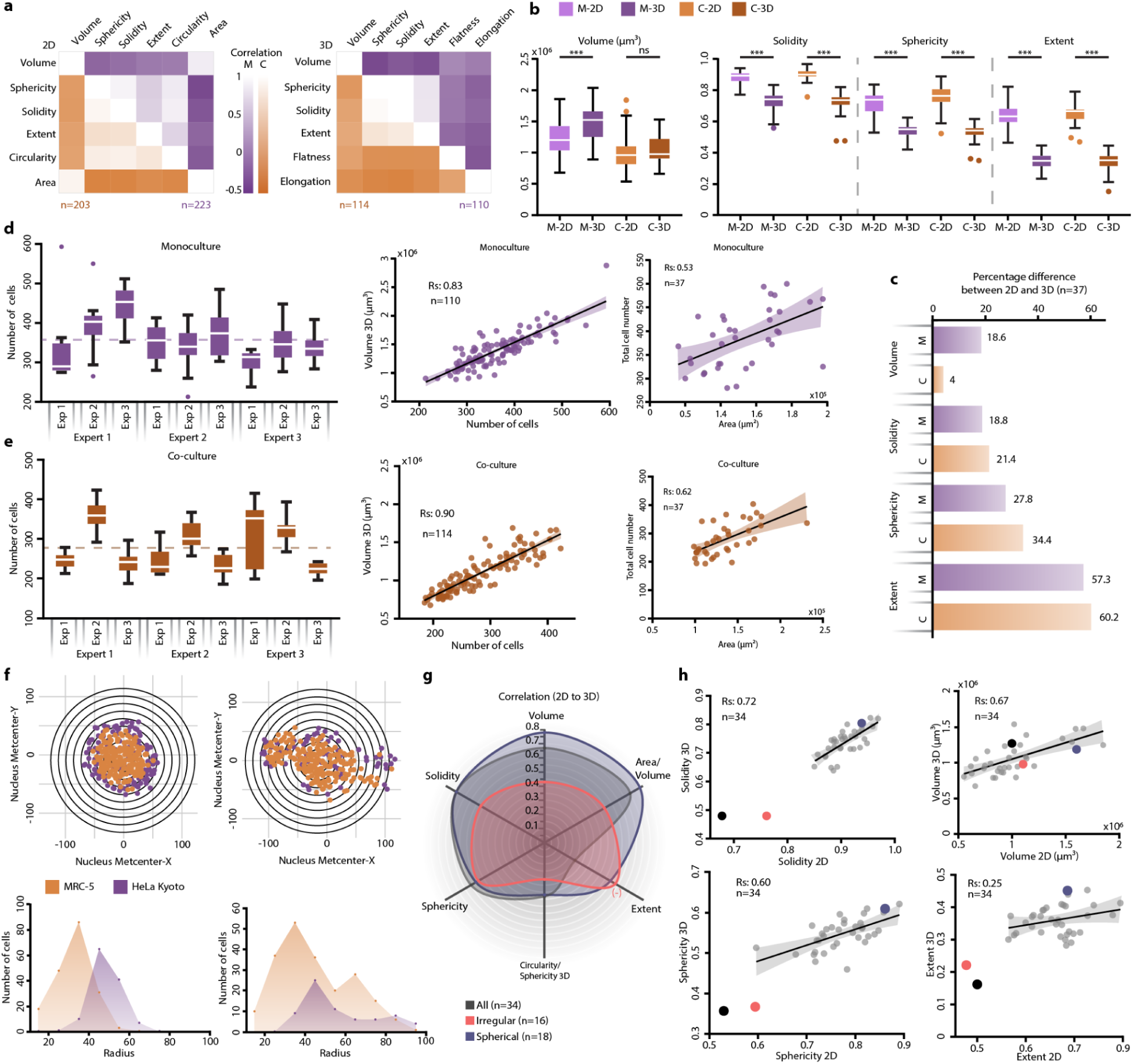
Comparison of 2D and 3D features. (**a**) Heatmap representation of correlations between features. Correlation between the selected features was tested using Spearman’s correlation method. Purple colour represents the monoculture dataset (n=223 for 2D and n=110 for 3D) and the orange colour represents the features collected from the co-culture dataset (n=203 for 2D and n=114 for 3D). (**b**) Boxplot representation of the 2D and 3D features. Thirty-seven spheroids were selected and verified that the 2D measurements correspond to the same objects in 3D. Features such as *Volume 3D* (µm^3^), *Solidity* (−), *Sphericity* (−), and *Extent* (−) were extracted from both the 2D and 3D images and used for comparison. Results are displayed as Monoculture 2D (light purple, M-2D), Monoculture 3D (dark purple, M-3D), Co-culture 2D (light orange, C-2D), and Co-culture 3D (dark orange, C-3D). The Kolmogorov-Smirnov test was used. *p ≤ 0.05; **p ≤ 0.01; ***p ≤ 0.001. (**c**) Percentage difference from the average values between the 2D and 3D features are shown. Purple colour represents the monoculture dataset, orange colour represents the co-culture dataset (n=37). (**d**) Boxplot representation of the number of nuclei in the monoculture dataset for each experiment (n≥7). Significant differences were not included. Correlation was measured between the number of nuclei, *Volume 3D*, and *Area*. (**e**) Boxplot representation of the number of nuclei in the co-culture dataset for each experiment (n≥7). Significant differences were not included. Correlation is measured between the number of nuclei, *Volume 3D*, and *Area*. (**f**) Vertical projection of a spherical and irregular spheroid that displays the number of HeLa Kyoto (purple) and MRC-5 (orange) cells. The center of each spheroid was defined using whole spheroid segmentation. Starting from the center position, spheres with increasing radius were used to create shells and count the number of objects in each section. Line plots show the number of cells per class based on the shell analysis. (**g**) Spider plot representation of the Spearman’s correlation between 2D and 3D features. Paired spheroids ranked by *Solidity 3D* (grey, n=34) are divided into a Spherical (dark blue, n=18) and Irregular (red, n=16) groups. (**h**) Correlation between 2D and 3D features (n=34). Previously displayed Spherical (dark blue) and Irregular (red) spheroid, along with the paired samples are displayed. Outliers (black and red) were not used for the evaluation. For the statistical analysis, a non-parametric Kruskal-Wallis test was conducted, followed by Dunn’s multiple comparison test. *p ≤ 0.05; **p ≤ 0.01; ***p ≤ 0.001.

To provide additional information relevant to phenotypic screening, we examined nucleus segmentation using the HCS system (**Fig. 3d,e**). For the monocultures, the average number of nuclei was 358, while the biggest spheroid (585) had 2.76 times more nuclei than the smallest (212 nuclei) (**Supplementary Fig. 4**). For the co-culture spheroids, the average number of nuclei was 280, and the biggest spheroid (423) had 2.32 times more nuclei than the smallest (185) (**Supplementary Fig. 5**). While both datasets showed a high positive correlation between the number of cells and the corresponding *Volume 3D*, only a moderate positive correlation was observed by changing volume to *Area* (**Fig. 3d,e**). No correlation was measured for *Solidity* and the cell number (**Supplementary Fig. 3d**). Further divergence between the cell number and *Area* was noted when the percentage difference was calculated for each experiment (**Supplementary Fig. 3c**). In both cases, co-culture spheroids reached a higher correlation (0.90 for *Volume 3D* and 0.62 for *Area*) than monocultures (0.83 for *Volume 3D* and 0.53 for *Area*).

Next, we selected a Spherical and an Irregular spheroid from co-cultures as representative cases (**Fig. 1n** and **Fig. 3f**). The spherical sample revealed that MRC-5 cells were grouped in the middle of the spheroid and surrounded by the HeLa Kyoto cells (**Supplementary Fig. 6a-c**). This structure, a fibroblast core covered by the tumour cells, was more common and spheroids with a more spherical shape showed similar distributions. Meanwhile, the structure of the irregular spheroid had multiple MRC-5 cores and HeLa Kyoto cells were more evenly distributed. The position of the cells within a spheroid proved to be a good indication of the structure, however, since the point of intersection is a characteristic feature due to size and the total number of cells.

Among the 37 spheroids, we created a ranking using *Solidity 3D* and divided the dataset into *Spherical* and *Irregular* groups to understand which morphology is more predictable based on the 2D features. *Volume*, *Circularity*/*Volume 3D*, and *Solidity* showed the highest positive correlations for the spherical group, 0.78, 0.78, and 0.72. *Sphericity* and *Circularity/Sphericity 3D* showed a moderate positive correlation (**Fig. 3g,h**). In general, the Spherical group always reached a higher correlation than the *Irregular* group (*i.e.* amorph spheroids), indicating that the more regular shape resulted in better predictability. Visualising spheroids by their shape (*Solidity*, *Sphericity*, *Extent*), two outliers (one of them is the Irregular spheroid) were distinguished and removed from the analysis (**Fig. 3h**). However, relying only on the size information, the removed outliers cannot be separated from the other samples.

Due to the nature of 3D-oids, comparing hundreds of samples may always show high heterogeneity. Utilizing 2D parameters while excluding 3D considerations allows faster and less demanding analysis resulting in overall greater similarity between samples. Perfectly spherical models are ideal for estimating 3D properties (*e.g. Volume 2D* and *Solidity*) since other 3D-oids may show irregular shapes that reduce predictability and comparability. For any 2D analysis, using at least 2 non-correlating features, such as size and shape descriptors, potentially helps to remove strong outliers. However, a 2D approach is insufficient to select the most similar spheroids in 3D.

### 3D single-cell analysis of co-culture spheroid model

A total of 114 HeLa Kyoto/MRC-5 spheroids were analysed at a single-cell level through segmentation and classification using a custom-developed analysis pipeline in *BIAS* (**Fig. 4a-d** and **Supplementary video 4**). Results on cell ratio showed that only 24% of the 114 spheroids predominantly consisted of HeLa Kyoto cells (hereafter called *H-M group)*, while 63% primarily contained MRC-5 cells (hereafter called *M-M group*) (**Fig. 4a**). Only 14 spheroids showed equal numbers of the 2 cell lines (hereafter called *E group*). The dataset included 32012 segmented objects where 18659 were classified as MRC-5 and 13353 HeLa Kyoto cells. By dividing spheroids based on their class (*H-M*, *E*, and *M-M groups*), higher total cell number was identified for the *M-M group* (**Fig. 4a**). Cytoplasm segmentation showed that the tumour cell line has significantly bigger volume, almost double the size of the fibroblast cells (**Fig. 4b**). Features extracted from the cellular microenvironment (*i.e*., neighbourhood features) further support the size differences since the distances between the nuclei and the cells are increased in favour of HeLa Kyoto cells. Considering the size difference between the cell lines, we wanted to test whether the different cell ratio changes the morphology of the spheroids. The *H-M group* showed significantly smaller spheroids and there was no significant difference for *Solidity* (**Fig. 4c and Supplementary Fig. 3e**). Although each spheroid is supposed to have a similar cell number, significant differences were measured concerning the number of cells for the different compositions (**Supplementary Fig. 3f**). Correlation between the total cell number and *Volume 3D* showed the highest correlation for the *E group* (Rs: 0.91), followed by the *M-M* (Rs: 0.86) and *H-M* (Rs: 0.79) (**Fig. 4d**).

**Fig. 4:**
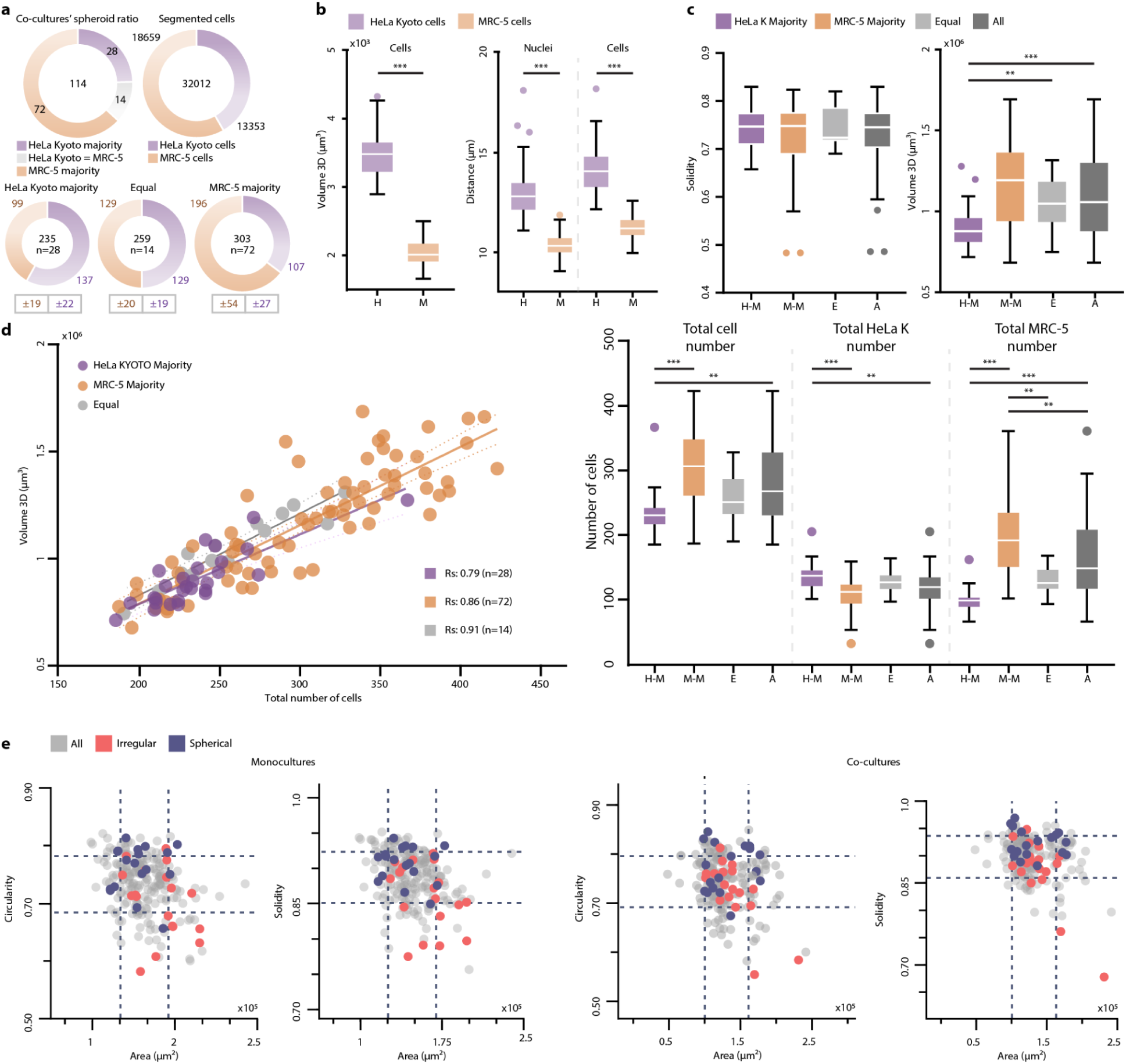
Single-cell analysis of co-culture spheroids. (**a**) Cell ratio for the 114 co-culture spheroids, showed 72 spheroids with MRC-5 cells in the majority (orange), 28 spheroids containing a majority of HeLa Kyoto (purple), and 14 spheroids with an equal number (grey). Ratio of classified objects with the exact number are displayed. The 114 spheroids were divided into 3 groups based on the cell type majority (*i.e*. HeLa Kyoto majority, Equal, MRC-5 majority). The size of the doughnut charts and the numbers in their centre represent the average cell numbers for each group. The average number of cells for each class is displayed with orange for HeLa Kyoto and purple for MRC-5. (**b**) Comparison of the two cell lines based on *Volume 3D* (µm^3^), the average inter-nuclear, and cell distance of the same class in µm. (**c**) Features (*Solidity*, *Volume 3D*, number of cells) collected from the spheroids were displayed based on the 3 groups. Dark grey colour represents the average value of the whole dataset. (**d**) Spearman’s correlation of the total number of cells and *Volume 3D* were visualised based on HeLa Kyoto majority (purple), MRC-5 majority (orange), and Equal (grey) groups separately. n=114. (**e**) 36 spheroids ranked by *Solidity 3D*, are separated into Spherical (dark blue) and Irregular (red) groups. These spheroids along with the total dataset (grey) were plotted in a scatter plot using only 2D features (*Circularity*, *Solidity*, and *Area*). Each circle represents a spheroid and dashed lines represent the standard deviation. n=223 and n=203 for the monoculture and co-culture dataset. For the statistical analysis, a non-parametric Kruskal-Wallis test was conducted, followed by Dunn’s multiple comparison test. When comparing only two groups, the Kolmogorov-Smirnov test was utilized. *p ≤ 0.05; **p ≤ 0.01; ***p ≤ 0.001.

As a final experiment, we were interested in whether examining 2D features alone is enough to select ideal spheroids. The 36 paired spheroid images ranked by *Solidity 3D* and separated into 2 groups (above) were used for this analysis. We plotted and highlighted *Spherical* and *Irregular* spheroids using the highest correlating 2D features (*Area*, *Solidity 2D*, and *Circularity)* for comparison. The scatter plot showed that the 2 groups are indistinguishable and data points cannot be visually separated. Relying on *Circularity*, spheroids from the Spherical group are outside of the standard deviation (**Fig. 4e**). Although *Solidity 2D* showed slightly better separation of the 2 groups compared to *Circularity*, the data points are intermixed and cannot be visually distinguished.

## Discussion

This study has developed, validated and presented HCS-3D*X*, a customisable 3D imaging and HCS system for analysing 3D-oids at a single-cell level. This innovative system addresses several challenges associated with 3D HCS, including the pre-selection, handling, imaging, and analysis of 3D-oids.

A critical aspect of this work was developing an HCS plate that can separate and allow fast, automated screening of multiple objects on the same plate, where the position of the samples is predefined. The plate is optimized to overcome time-consuming sample preparation processes. Nevertheless, it is highly adaptable and customisable, allowing modifications to be made within hours to suit individual needs. The plate was developed and tested on an upright light-sheet microscope to achieve single-cell resolution in spheroids. However, any type of 3D cell culture can be screened.

We proved that the designed multiwell plate doubles screening speed and provides high penetration depth, 250 or even 350 µm for optically cleared tumour spheroids^28,29^ using a Leica SP8 TCS DLS microscope. Since FEP foil allows gas exchange and LSFM offers reduced phototoxicity and increased imaging speed, we found this combination the most ideal for long-term live imaging of the specimen. Furthermore, compatibility with a multi-pipette or a pipetting robot, *SpheroidPicker*, allows AI-based pre-selection and an automated transfer process (**Supplementary Note 4**).

Several previous publications report that pre-selecting spheroids is crucial in many aspects but for many reasons, reproducibility is the most essential^15,18,30–33^.

Our understanding of a 3D-oid depends on the extracted features. However, we demonstrated that relying only on 2D features limits the analysis, leading to an incomplete representation of the data. We compared features extracted from brightfield (2D) and fluorescence (3D) images of spheroid models to quantify and understand the importance of losing one dimension. Considering image-based features, certain 2D parameters are more important than others when pre-selection is an option. Most publications describe 3D-oids using *Diameter*, *Perimeter*, *Area*, *Volume*, *Circularity*, *Sphericity*, and *Convexity*, estimated using images acquired at various magnifications^31,32,34^. While the choice of different objectives depends on many aspects (size, speed, working distance, etc.), we wanted to see whether the variability causes a problem for comparing different studies. Comparing 4 different objectives ranging from 2.5x-20x, the average differences were less than 5%, and significant differences were measured between the 2.5x, the 5x and 10x objectives. Among the tested features, we observed that *Perimeter*, *Convexity*, *Sphericity*, and *Circularity* showed significant differences. The higher magnification images resulted in more precise contours that follow the edge of the objects accurately, while annotations on images taken with lower magnification objectives were less accurate. Thus, annotating images taken at different magnifications caused discrepancies for *Perimeter* and for all the features that rely on it (*e.g. Circularity, Sphericity*, and *Convexity*). In addition to the objective’s physical parameters (such as magnification, numerical aperture, and spherical aberration), the lower image resolution and the accuracy of the annotator caused the general inconsistency between the objectives.

We suggest a 10x objective provides a fair compromise for general feature reliability if 20x is unavailable or has an excessively narrow field of view, even though the 2.5x or 5x objectives would not distort the entire analysis. In addition, available AI-based segmentation models can improve the consistency of object detection and reduce time-consuming annotation tasks.

As 3D-oids pre-selection is often recommended, it is important to decide which features are ideal for selection. By discarding one dimension, using *Diameter* instead of *Area*, the spheroids are numerically more similar to each other. In addition, *Diameter* and *Circularity* showed moderate negative correlations (Rs: −0.69 and −0.54), indicating that smaller spheroids were more circular and more spheroids were found to be ideal. Such correlation is not general since different 3D models may show opposite results^35^. Plotting spheroids based on *Area* and *Circularity* showed no correlation and less spheroids as ideal. *Volume* estimated using 2D features resulted in slightly smaller values but proved to be more accurate when spheroids were more rounded. Other 2D shape descriptors like *Solidity*, *Sphericity*, and *Extent* showed significantly lower values compared to the measured 3D values. By measuring exactly the same spheroids both in 2D and 3D, *Volume*, *Area/Volume 3D*, and *Solidity* showed the highest correlations (*i.e*. 0.78, 0.78, and 0.72, respectively). However, highest correlation is only possible for the most spherical samples and irregular objects decrease the predictability. Correlation between the number of segmented objects to *Volume 3D* showed a strong positive correlation (0.83 and 0.90), whilst *Area* showed weaker correlation (0.53 and 0.62). In conclusion, AI-based nucleus segmentation accurately represents spheroids, whilst estimating the number of cells based on the *Area* of the sample is not accurate.

Shape and size features showed moderate or no correlation, thus the number of outliers cannot be distinguished using only one feature. Comparing spheroids in 2D and 3D showed that describing the population of spheroids requires at least two parameters relevant to the size and the shape. While pre-selection certainly helps to remove strong outliers, it does not select the most similar spheroids in 3D.

The heterogeneity of spheroids is influenced by the type of cells, the environment, and the formation method, indicating the complexity of generating 3D cell cultures^13^. To measure the variability of the spheroid models, 3 experts repeated the same experiments 3 times generating 426 monoculture (223) and co-culture (203) spheroids that were analysed in 2D and 3D. Despite using identical equipment and working in the same environment, the comparison of the manual plating revealed great heterogeneity in the size and morphology of spheroids. Furthermore, significant differences were measured by comparing the performance of the various experts. By generating spheroids from a single cell line, we demonstrated that a higher number of replicates and a proper 2D feature quantification combined with pre-selection is recommended.

A particularly insightful evaluation showed a significant difference for the size but no difference for circularity, indicating that spheroid shape was unaffected by size changes. As expected, cell type and ratio had a major impact on spheroid formation. Although co-cultures were seeded with twice as many cells, those spheroids were significantly smaller. Thus, the cell type and ratio has a significant effect on spheroid formation. Interestingly, co-culture spheroids showed higher positive correlation for the size and number of cells than monocultures, even though the 2 different cell types showed unequal sizes in spheroids.

Increasing the complexity of the biological model introduces more bias into the experiment, potentially leading to incorrect conclusions. Apart from size, shape, and number of cells, the cell ratio and structural information are also needed to describe a multi-cell line spheroid. Achieving single-cell resolution and applying 3D analysis (including segmentation, classification, and feature extraction), 114 co-culture spheroids were analysed and compared at a cellular level. After evaluating the individual cells composing the spheroids, we observed that spheroids can be categorised into 3 groups according to the various cell ratios and total cell numbers. Although the groups showed significant differences in size, there was no inequality in shape. The total number of cells revealed that spheroids with more cells contained more MRC-5 cells than HeLa Kyoto.

In this large-scale, single-cell analysis experiment, we demonstrated that complex models relying on interactions between various cell types result in high variety. However, the consistency of morphologically similar spheroids also varies with the expert conducting the experiment.

However, pre-selection methods for 2D spheroids did not bring the promised impact in this field, it is still essential to reduce the model heterogeneity and the high variability resulting from the generation process^34^. It should be pointed out that 2D pre-selection applied to more complex models may eliminate one specific type of spheroid with different cell ratio rather than selecting the most similar ones. Therefore, size and structural differences between 3D-oid models emphasises the need for a single-cell analysis pipeline to avoid incorrect data interpretations and false conclusions.

Over the years, only a few methods were published for 3D-HCS. Widefield and confocal fluorescence systems are more frequently applied since such systems are compatible with standard plate formats, thus more commonly used for screening. Widefield fluorescence imaging offers high screening speed and compatibility with microfluidic systems, but it has low penetration depth and only allows limited feature extraction^36,37^. On the other hand, confocal fluorescence microscopy systems have subcellular resolution but with a much slower image acquisition and higher phototoxicity^12,38–40^. In addition, due to the limited light penetration, HCS methods are usually demonstrated on smaller 3D-oids using standard but more expensive imaging plates^41^.

HCS systems designed for LSFM overcome most of the disadvantages offering single-cell resolution images with fast image acquisition, however, complex sample preparation and the usage of special plates is required^24,42–44^. Furthermore, such systems are usually specific to the imaging setup resulting in limitations (i.e. size or type specific imaging plate, manufacturing price and protocol of the plate, modifications of the microscope, availability and complexity of code).

Working with 3D-oids has resulted in significant variability, thus the developments aimed at evaluating the samples must also be flexible. Imaging and image analysis systems are needed that ensure the interchangeability of individual components, transparency in data reporting, and accessibility of the code. A comprehensive method review is presented in **Supplementary Table 2**.

In this work, we developed HCS-3D*X,* a versatile HCS system which comprises all the steps needed to evaluate 3D cell cultures from the beginning to the end at a single-cell level. By integrating AI-driven solutions and advanced imaging techniques such as LSFM, HCS-3D*X* enables precise, automated, and high-throughput analyses at a single-cell level. Accordingly, it represents a substantial leap forward in 3D HCS, offering a comprehensive solution that bridges the gap between traditional 2D models and complex *in vivo* environments. Its application will enhance the accuracy and efficiency of drug screening processes, support personalised medicine approaches, and facilitate more detailed investigations into cellular behaviour within 3D structures. In conclusion, its robustness and versatility make HCS-3D*X* a valuable tool for researchers and clinicians, paving the way for future innovations in biomedical research.

## Materials and Methods

### Spheroid models

Spheroid monocultures were generated using HeLa Kyoto EGFP-alpha-tubulin/H2B-mCherry cervical cancer cells (Cell Lines Service, Eppelheim, Germany). Cells were maintained in a HeLa Kyoto medium consisting of DMEM (Lonza, Basel, Switzerland), 10% Fetal Bovine Serum (FBS, Euroclone, Milan, Italy), 2 mM L-glutamine (Lonza), 0.5 mg/ml G418 (Gibco, Montana, United States), and 0.5 µg/ml puromycin (Sigma, Kanagawa, Japan). To generate uniform spheroids, 100 cells were seeded into each well in U-bottom cell-repellent 384-well plates (Greiner Bio-One, Kremsmünster, Austria) for 48 h at 37 °C and 5% CO_2_. After 48 h, spheroids were collected and then washed 3 times with Dulbecco’s Phosphate Buffered Saline (DPBS), and fixed with 4% Paraformaldehyde (PFA) for 60 min. Spheroids were washed with DPBS 3 times and stored at 4°C in DPBS until imaging. Before imaging, spheroids were incubated in 0.1% Triton X-100 overnight at room temperature and washed 3 times with DPBS. For actin labelling, spheroids were stained with 1:200 Flash Phalloidin NIR 647 (Biolegend, San Diego, California) for 60 min. Before the imaging, spheroids were washed with DPBS 3 times. T-47D spheroids used for the image quality test were generated according to previous publication^29^.

Co-culture spheroids were generated using the same HeLa Kyoto cells and MRC-5 fibroblasts (American Type Culture Collection - ATCC). The manufacturer’s instructions were followed for the maintenance of the cell cultures. To generate spheroids, a co-culture medium consisting of DMEM, 10% FBS, 1% L-glutamine (2 mM), and 1% Penicillin-Streptomycin-Amphotericin B mixture (all from Lonza) was used. 40 HeLa Kyoto cells were seeded into each well in U-bottom cell-repellent 384-well plates at 37 °C and 5% CO_2_. After 24 h of incubation, 160 MRC-5 cells per well were added onto the HeLa Kyoto cells and the co-cultures were incubated for one more day. After 24 h, the co-culture spheroids were collected from each well and washed 3 times with DPBS. 4% PFA was used for 60 min to fix the samples, then washed again with DPBS 3 times and stored at 4°C in DPBS until imaging. Before imaging, spheroids were incubated in 0.1% Triton X-100 and 1 µg/ml DAPI overnight at room temperature and washed 3 times with DPBS. Spheroids were stained with 1:200 Flash Phalloidin NIR 647 for 60 min. Finally, spheroids were washed with DPBS 3 times.

Hydrogel-based multicellular human tumour models were generated by co-culturing CellTracker Orange CMTMR (Invitrogen)-stained T-47D ductal carcinoma and stroma cells, i.e., CellTracker Deep Red-stained MRC-5 fibroblasts, and CellTracker Green CMFDA-stained EA.hy926 endothelial cells in hydrogel matrix (TrueGel3D, Sigma-Aldrich) directly in the HCS plate.

### HCS foil multiwell plate

A customisable 3D imaging multiwell plate was designed for the screening of 3D-oids at a single-cell level. The plate includes: (*I*) a 3D printed base element that retains the detection fluid; (*II*) a FEP foil for separating the position of the samples; (*III*) an insert element to secure the samples’ position within the foil; and (*IV*) a 3D printed grid element to fix the FEP foil position (**Fig. 3**). This designed plate is suitable for examining a large number of samples with an LSFM, with a single sample in each cuvette. The main parts of the plate are briefly discussed in the following sections, whilst detailed information about the dimensions and assembly are available in **Supplementary Note 1**.

FEP foils are fully transparent, with a refractive index of 1.341-1.347. FEP is chemically inert and resistant to organic solvents, acids, and bases, similar to the majority of fluorocarbon plastics. Additionally, the material satisfies FDA (21CFR.177.1550) and EU (2002/72/EC) requirements^20^. FEP-films (Holscot Europe, Netherlands) with a 50 µm thickness were cut into 15 x 15 cm pieces and cleaned with 70% alcohol. Each foil was placed and clamped into the frame of a JT-18 vacuum-forming machine (Yuyao Jintai Machine Company, China). To get small and uniform shapes, the heater needs to raise the temperature to a level near the glass transition temperature of the FEP-foil (260–280°C). Once the heater reaches the desired temperature, the positive mould is quickly placed onto the vacuum-forming machine, and the foil is quickly pressed onto the mould whilst the vacuum suction is switched on. The FEP foil that has been extruded is then carefully taken out of the mould and cleaned with an isopropyl alcohol bath for 5 min to remove the excess resin components.

Using *Blender 3.0*, positive moulds for the spikes were created. Each mould was designed to shape the FEP foil in order to fit perfectly into the base element. For printing, the 3D models were exported to a stereolithographic file format (.stl) and then imported into *PrusaSlicer v 2.6.x* (Prusa Research, Czech Republic). Each heat-resistant mould was printed with the Prusa SL1 printer, a stereolithography (SLA) resin-based 3D printer that forms objects layer by layer using a liquid resin that is ultraviolet (UV)-cured. This printer provides high-resolution printing capabilities enabling the production of intricate and detailed objects with smooth surfaces. The mechanical and thermal characteristics of heat-resistant resin enable it to endure the vacuum-forming process. The positive moulds were UV-cured and washed with isopropanol to remove excess resin from the surface. Then the moulds were examined under a stereomicroscope and cleaned by submerging them in an ultrasonic bath before being used.

The base components of the plates were printed with the Prusa i3 MK3S+ 3D printer with PETG (3DP-PETG1.75-01-BK; Gembird, Shenzhen, China) in two different sizes. The model was sliced with *PrusaSlicer*. 3D printed with default Prusa PETG filament profile and with “0.2 Quality print settings” using a 0.4 mm nozzle. The bigger version is 85 / 127 / 15.4 mm (average size of a 96-well plate) which is suitable for screening 56 samples, while the smaller version 85 / 73.8 / 15.4 mm was designed only for 28 samples. To create a waterproof base for the samples, the 24 / 64 mm cover glasses (Menzel Gläser, Germany) were secured with silicon grease (ThermoFisher, USA) to the bottom of the bases. The cover glass provides a transparent bottom for any inverted microscopy setup while allowing the freedom of using dipping objectives.

A grid element was 3D printed using a Prusa i3 MK3S+ printer with PETG to prevent the samples from shifting after being filled with mounting media. The grid element can be slid into predefined spaces, securing the foil to the bottom of the plate.

### Microscopes

For brightfield imaging, the fixed monoculture and co-culture spheroids were placed into a 35/10 mm cell culture dish with a glass bottom (627965, Cellview, Austria) in PBS. Brightfield images were taken with the Leica SP8 TCS using 4 different objectives: 2.5x/0.07, 5x/0.15, 10x/0.32, and 20x/0.4 (**Supplementary Table 3**).

The HCS foil multiwell plate was validated on the Leica TCS SP8 Digital LightSheet (DLS) microscope, exploiting a standard 96-well plate insert. A 25x/0.95 detection objective with a mounted 2.5 mm mirror device was used to capture fluorescence DLS images, illuminated by the 2.5x/0.07 objective. Images with a resolution of 2048×2048 pixels and a pixel size of 0.144 µm were captured with the sCMOS DFC9000 Leica camera. A 2 µm gap distance between the images was used in each z-stack. dH_2_O mounting medium was used for every spheroid. 3 different tests were performed changing laser and laser intensity (**Supplementary Fig. 2** and **Fig. 4**). Firstly, a laser with a wavelength of 638 nm and an exposure time of 200 ms with 20% laser intensity (maximum laser intensity 350 mW) was used. Then, laser intensity was manually adjusted for each channel (*i.e*. 405, 488, 552, and 638 nm wavelength).

### Image analysis

Brightfield images were annotated using the *AnaSP* software^45^ to obtain binary masks and *Volume* features. Original brightfield images and corresponding masks were imported into *BIAS* and 2D features were exported^26^. Image quality was evaluated using intensity variance, a metric implemented in *Spheroid Quality Measurement* (SQM), an open-source *ImageJ*/*Fiji* plugin^28^. All the 3D fluorescent image stacks were directly analysed in *BIAS* using algorithms based on U-Net^46^ (for whole spheroid segmentation) and *StarDist*^48^ (nuclei segmentation) for segmentation and multilayer perceptron (MLP) for classification (**Supplementary note 2 and 3**).

### Statistical analysis

Statistical analyses were conducted using *GraphPad Prism 8* software. The Kolmogorov-Smirnov test was applied to assess normal distribution. For analysing the results of the 2D features, the non-parametric Kruskal-Wallis test followed by Dunn’s multiple comparisons was used. The significance level was set at α = 0.05 with a 95% confidence interval, and p-values were adjusted for multiple comparisons.

## Acknowledgements

The authors would like to thank Lilla Pintér (HUN-REN BRC, Szeged, Hungary) for her technical support. We acknowledge support from the Lendület BIOMAG grant (no. 2018–342), TKP2021-EGA09, HUNRENTECH (TECH-2024-34), Horizon-BIALYMPH, Horizon-SYMMETRY, Horizon-SWEEPICS, H2020-Fair-CHARM, HAS-NAP3, the HUNTER-Excellence 2024, grant from OTKA-SNN no. 139455/ARRS and OTKA-Excellence 2025, the FIMM High Content Imaging and Analysis Unit (FIMM-HCA; HiLIFE-HELMI), and Finnish Cancer Society. F.P. acknowledges support from the MAECI Science and Technology Cooperation Italy-South Korea Grant Years 2023–2025 by the Italian Ministry of Foreign Affairs and International Cooperation (CUP project: J53C23000300003); M.H. from the ÚNKP-23-4 -SZTE-639 New National Excellence Program of the Ministry for Culture and Innovation from the source of the National Research, Development and Innovation Fund.

## Author contributions

Conceptualisation: A.D., M.H., F.Pi., P.H.;

Methodology: A.D., T.T., I.G., B.S., M.H.;

Software: F.K., A.K.;

Validation: A.D., T.T., I.G., N.H.;

Formal analysis: A.D., M.H., F.Pi., P.H.;

Investigation: A.D., T.T., I.G., B.S., N.H., F.Pa.;

Resources: A.D., I.G., F.K., A.K., M.H., K.B., F.Pi., P.H.;

Data curation: A.D., T.T., I.G., B.S., M.H.;

Writing-original draft preparation: A.D., M.H., F.Pi.;

Writing-review and editing: T.T., I.G., B.S., N.H., F.K., A.K., K.B., F.Pa., P.H.;

Visualisation: A.D., B.S.;

Supervision: K.B., F.Pa., P.H.;

Project administration: A.D., K.B., F.Pi., P.H.;

Funding acquisition: K.B., F.Pi., P.H.;

All authors read and approved the final version of the manuscript.

## Ethics declarations

## Data and materials availability

All data needed to evaluate the conclusions in the paper are present in the paper and/or the Supplementary Materials. For further details and data referenced in this article, please visit the links provided below:

Monoculture and co-culture spheroid image dataset: https://doi.org/10.6084/m9.figshare.c.7658858

Optically cleared spheroids: https://doi.org/10.6084/m9.figshare.c.5051999.v1

HCS of tumour-stroma spheroid multicultures: https://doi.org/10.6084/m9.figshare.c.7357135

Annotated 3D image dataset: https://doi.org/10.6084/m9.figshare.c.7020531.v1

SpheroidPicker:

● Source code: https://github.com/grexai/SpheroidPicker
● The model files (3D printed elements): https://doi.org/10.5281/zenodo.14679243
● Original and the improved segmentation models: https://doi.org/10.5281/zenodo.14675683
● Annotated dataset: https://doi.org/10.5281/zenodo.14679303

BIAS software: https://single-cell-technologies.com/

Additional data related to this paper may be requested from the corresponding author.

## Supplementary Materials

**Supplementary video 1: Concept of the HCS-3D*X* workflow.** Demonstrative video of the individual steps using each component of the HCS-3DX pipeline featuring the 3D-oid pre-selection, treatment, and imaging.

**Supplementary video 2: Optically cleared tumour spheroid.** Example video of an optically cleared T-47D spheroid labelled with DRAQ5 for nuclei (red) and Phalloidin 488 for actin (cyan). The spheroids were optically cleared using a sucrose-based clearing protocol. A total of 5,184 nuclei were segmented with the BIAS software. Scale bar represents 100 µm.

**Supplementary video 3: Live-cell imaging of cells embedded in hydrogel.** By co-culturing T-47D breast cancer cells with MRC-5 fibroblasts and EA.hy926 endothelial cells, a hydrogel-based tumour model was created to track the tumour microenvironment in real time. First, the whole region of the FEP-foil was imaged with a lower magnification objective (10x detection objective; z-step: 5 µm; ∼300 µm in z). Then specific regions were selected and imaged with a higher magnification objective (25x detection objective; z-step: 2 µm; ∼200 µm in z) using the LSFM. CellTracker dyes (orange, deep red, and green) are used to detect cells. The tumour, endothelial, and fibroblast cells are marked by red, green, and white labels, respectively. Nuclei of the cells were labeled with Hoechst 33342. For the long-term experiment, images were taken every 30 minutes for 30 hours using the HCS plate. Maximum intensity projection video was exported using the LAS X software (Leica). Scale bar represents 20 µm.

**Supplementary video 4: Spheroid analysis using BIAS.** Randomly selected co-culture spheroids were directly imported into BIAS for image analysis. The video demonstrates image analysis steps, such as nucleus segmentation, whole spheroid segmentation, and classification. Certain segments of the video have been accelerated and do not reflect real-time speed.

**Supplementary Table 1.** Number of publications spanning 2010 to 2024 including different queries based on PubMed (Public Medline). Full queries are available as an attachment. *Query 1*: Number of publications related to spheroids, organoids, tumoroids, tumouroids, and assembloids. *Query 2*: Number of publications related to spheroids, organoids, tumoroids, tumouroids, and assembloids, combined with various microscopy techniques like confocal, light-sheet, multiphoton, brightfield, and phase contrast. *Query 3*: Number of publications related to high content or high throughput screening, analysis, assays, systems, imaging, techniques, or pipelines. *Query 4*: Number of publications related to high content or high throughput screening, analysis, assays, systems, imaging, techniques, or pipelines, combined with various microscopy techniques like confocal, light-sheet, multiphoton, brightfield, and phase contrast. *Query 5*: Number of publications related to spheroids, organoids, tumoroids, tumouroids, and assembloids, combined with high content or high throughput screening, analysis, assays, systems, imaging, techniques, or pipelines, and various microscopy techniques like confocal, light-sheet, multiphoton, brightfield, and phase contrast. *Query 6*: Number of publications related to spheroids, organoids, tumoroids, tumouroids, and assembloids, combined with high content or high throughput screening, analysis, assays, systems, imaging, techniques, or pipelines, and specific microscopy techniques like confocal, light-sheet, or multiphoton, specifically focused on 3D.

| Year | *Query 1* | *Query 2* | *Query 3* | *Query 4* | *Query 5* | *Query 6* |
| --- | --- | --- | --- | --- | --- | --- |
| 2010 | 561 | 25 | 5153 | 53 | 3 | 1 |
| 2011 | 650 | 41 | 6145 | 69 | 1 | 1 |
| 2012 | 855 | 44 | 7334 | 85 | 1 | 1 |
| 2013 | 1072 | 55 | 8474 | 76 | 2 | 2 |
| 2014 | 1127 | 56 | 9554 | 84 | 5 | 4 |
| 2015 | 1366 | 78 | 11014 | 125 | 6 | 3 |
| 2016 | 1713 | 71 | 11987 | 110 | 5 | 4 |
| 2017 | 2135 | 102 | 12067 | 123 | 8 | 2 |
| 2018 | 2541 | 98 | 12934 | 127 | 14 | 9 |
| 2019 | 2948 | 116 | 13930 | 139 | 9 | 6 |
| 2020 | 3953 | 128 | 14559 | 125 | 10 | 7 |
| 2021 | 4801 | 137 | 15427 | 143 | 11 | 7 |
| 2022 | 4705 | 130 | 13672 | 131 | 21 | 14 |
| 2023 | 5114 | 136 | 12534 | 153 | 17 | 9 |
| 2024 | 5911 | 142 | 13768 | 147 | 9 | 6 |
| SUM | 39452 | 1359 | 168552 | 1690 | 122 | 76 |

**Supplementary Table 2.** Overview of HCS systems designed for 3D-oids. The columns in the table represent the following categories: (1) date of the publication; (2) automatic pre-selection and transferring of 3D-oids into imaging plates; (3) compatibility with automated liquid handling robots for fluid exchange; (4) utilization of specialized sample holders designed for imaging of 3D-oids; (5) type of the microscope used for the screening; (6) imaging technology; (7) type of the images that was used for analysis; (8) genuine utilisation of 3D information at the cellular level; (9) references (*Ref*). Abbreviations: WFM - Widefield fluorescence microscopy; CFM - Confocal fluorescence microscopy; LSFM - Light-sheet fluorescence microscopy, NA - Not available.

| # | Publication date | 3D-oids handling | Compatibility with pipetting robots | Special imaging plate included | Microscope | Technology | Image type | Single-cell resolution analysis | Ref |
| --- | --- | --- | --- | --- | --- | --- | --- | --- | --- |
| 1 | 2017 | no | yes | no | Zeiss Observer Z1 | WFM | 2D | no | 36 |
| 2 | 2018 | no | no | yes | ASFA™ Scanner ST, Medical & Bio Device, South Korea | WFM | 2D | no | 37 |
| 3 | 2019 | no | yes | no | Perkin Elmer Opera Phenix screening | CFM | 3D | yes | 39 |
| 4 | 2020 | no | no | yes | Perkin Elmer Opera Phenix screening | CFM | 2D | no | 40 |
| 5 | 2022 | no | no | yes | Zeiss Laser scanning microscopy 880 | CFM | 3D | no | 38 |
| 6 | 2024 | no | NA | NA | Leica TCS SP8 Confocal | CFM | 3D | yes | 47 |
| 7 | 2024 | no | yes | no | Perkin Elmer Opera Phenix screening | CFM | 2D | no | 41 |
| 8 | 2024 | no | yes | yes | Perkin Elmer Opera Phenix screening | CFM | 3D | yes | 12 |
| 9 | 2020 | no | yes | yes | Dual-view inverted selective plane illumination microscope | LSFM | 3D | yes | 42 |
| 10 | 2022 | no | no | yes | LS1-Live dual illumination and inverted detection microscope | LSFM | 3D | yes | 24 |
| 11 | 2022 | no | no | yes | Nikon Eclipse modified to LS | LSFM | 3D | yes | 43 |
| 12 | 2024 | no | no | yes | Leica TCS SP8 DLS | LSFM | 3D | no | 44 |
| 13 | 2025 | yes | yes | yes | Leica TCS SP8 DLS | LSFM | 3D | yes | OUR S |

**Supplementary Table 3.** Microscope settings and objectives used for imaging. Abbreviations: NA - numerical aperture, D - dry, W - water immersion.

| # | Microscope | Type | Objectives | NA | Pixel resolution | µm resolution | Coefficient | TwinFlect mirror |
| --- | --- | --- | --- | --- | --- | --- | --- | --- |
| 1 | Leica SP8 TCS | Detection | 2.5x | 0.07 D | 1392 x 1040 | 3588.78 x 2680.62 | 2.57783289 | - |
| 2 | Leica SP8 TCS | Detection | 5x | 0.15 D | 1392 x 1040 | 1794.39 x 1340.31 | 1.28891644 | - |
| 3 | Leica SP8 TCS | Detection | 10x | 0.32 D | 1392 x 1040 | 897.2 x 670.16 | 0.64446242 | - |
| 4 | Leica SP8 TCS | Detection | 20x | 0.4 D | 1392 x 1040 | 448.6 x 335.08 | 0.32223121 | - |
| 5 | Leica SP8 DLS | Detection | 10x | 0.3 W | 2048 x 2048 | 735.76 x 735.76 | 0.35925781 | 5 mm |
| 6 | Leica SP8 DLS | Detection | 25x | 0.95 W | 2048 x 2048 | 294.3 x 294.3 | 0.14370117 | 2.5 mm |
| 7 | Leica SP8 DLS | Illumination | 2.5x | 0.07 D | - | - | - | - |
| 8 | Leica SP8 DLS | Illumination | 5x | 0.15 D | - | - | - | - |

**Supplementary Table 4.** Relative differences between the various settings compared to the 20x objective. Spheroids were imaged with the Leica SP8 TCS microscope using 2.5x, 5x, 10x, and 20x objectives. The 2D morphological characteristics of 50 spheroids were analysed using BIAS and AnaSP. Relative differences were then calculated relative to the 20x objective for features such as *Diameter*, *Perimeter*, *Area*, *Volume*, *Circularity*, *Sphericity*, and *Convexity*. The average values for each objective are displayed as a percentage, n=50.

| # | Features | 2.5x objective | 5x objective | 10x objective |
| --- | --- | --- | --- | --- |
| 1 | *Diameter* | 1.59 | 1.42 | 1.15 |
| 2 | *Perimeter* | 4.15 | 2.06 | 1.70 |
| 3 | *Area* | 2.46 | 2.29 | 2.65 |
| 4 | *Volume* | 6.43 | 5.44 | 5.48 |
| 5 | *Circularity* | 3.57 | 1.32 | 1.18 |
| 6 | *Sphericity* | 4.81 | 1.90 | 2.10 |
| 7 | *Convexity* | 3.18 | 1.70 | 1.78 |
|  | Average | 3.74 | 2.30 | 2.29 |

**Supplementary Fig. 1.**
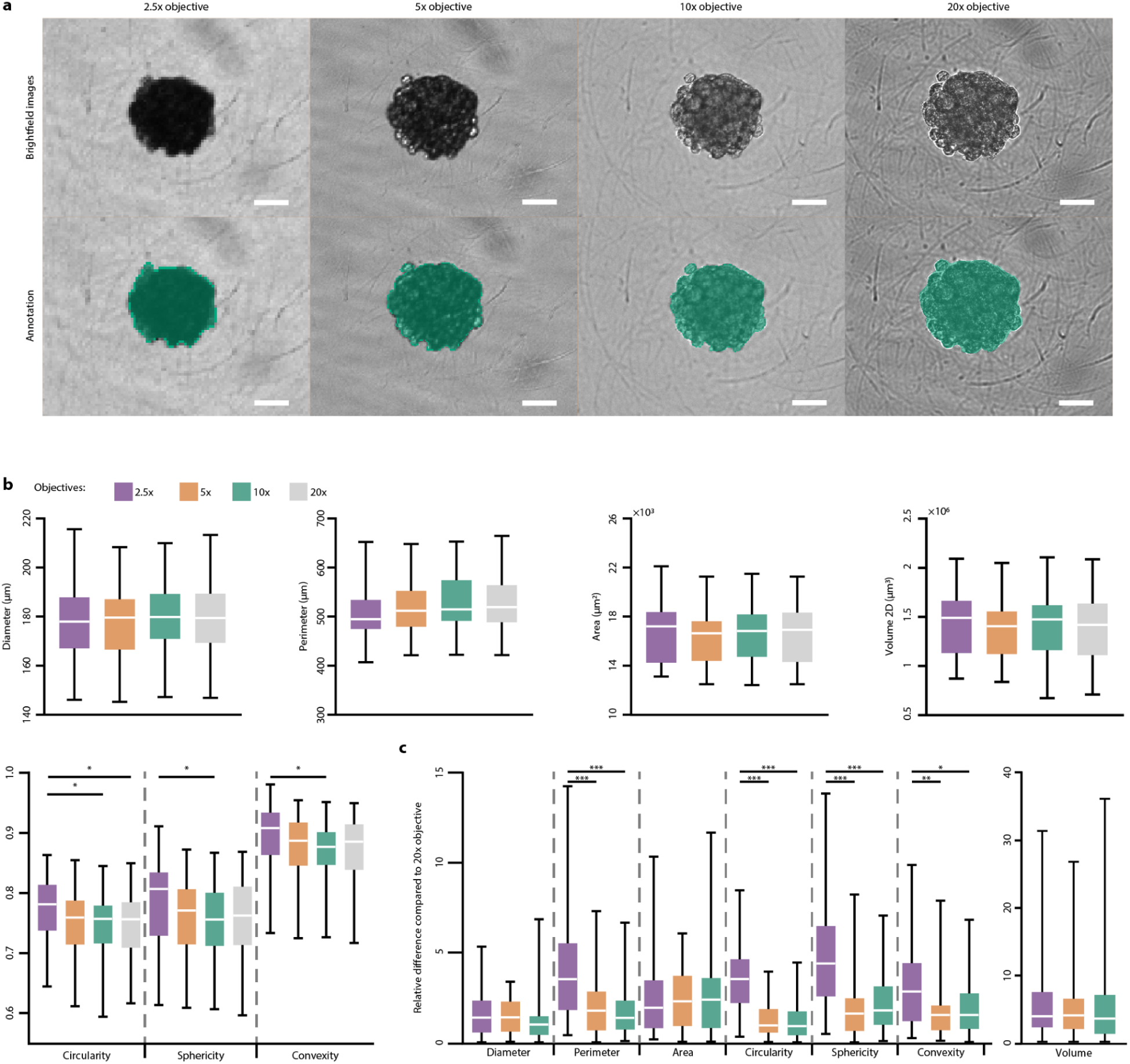
(**a**) Representative images of spheroids acquired with the 2.5x, 5x, 10x, and 20x objectives. Corresponding annotations for each sample are displayed below in green. Different zoom levels were used to enlarge spheroids to a similar size. The scale bar represents 50 µm. (**b**) Boxplot analysis of various 2D morphological features of 50 spheroids imaged by the Leica SP8 TCS microscope with 2.5x (purple), 5x (orange), 10x (green), and 20x (grey) objectives. Brightfield images of spheroids were manually annotated and 2D morphological features were extracted by using *BIAS* and *AnaSP*. Boxplot analysis of *Diameter* (µm), *Perimeter* (µm), *Area* (µm^2^), *Volume* (µm^3^), *Circularity* (−), *Sphericity* (−), and *Convexity* (−). (**c**) Figures represent the relative difference of the same features compared to the 20x objective (n=50). Each statistical test was carried out by features separately. *p ≤ 0.05; **p ≤ 0.01; ***p ≤ 0.001.

**Supplementary Fig. 2.**
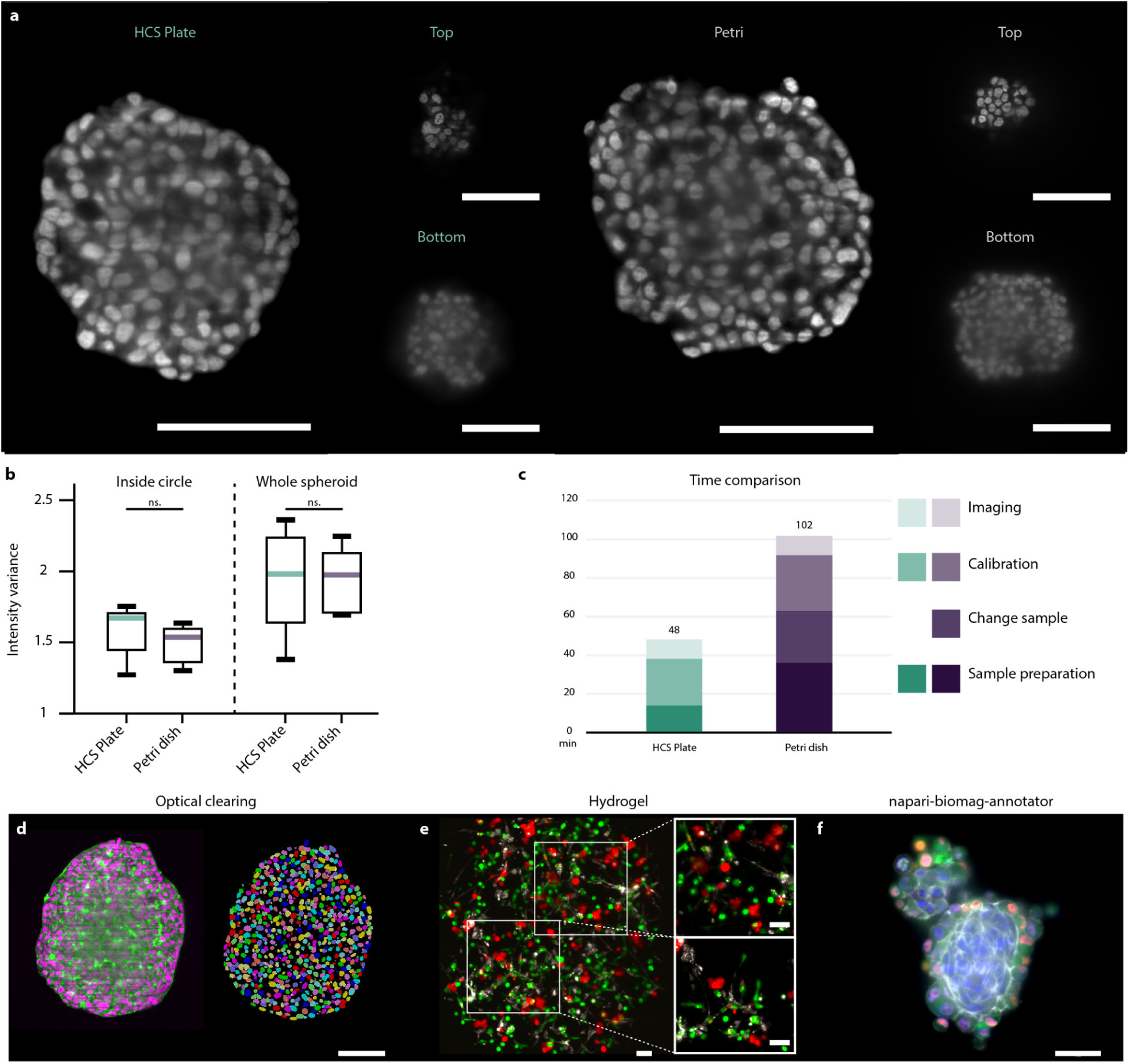
Qualitative comparison of the HCS plate and the traditional imaging method where the samples were placed into Petri dishes for LSFM. (**a**) Fluorescent images of T-47D spheroids^29^ stained with DRAQ5 were used as a reference to compare image quality. Randomly selected spheroids screened in the HCS plate (left image) and in Petri dish (right image) were visualized where images were selected from the top, middle, and bottom regions. Scale bars represent 100 µm. (**b**) To characterise the general blurriness of the whole z-stack, the intensity variance metric was used both for the inside circle and for the whole spheroid. Results are shown in a boxplot, showing that no significant differences were measured between the HCS plate and the Petri. n=5 for each group. (**c**) Preparation and screening times were measured for both techniques and displayed on a stacked bar plot. Starting from the darkest colour the 4 categories are Sample preparation, Change sample (this category does not apply to the HCS plate), Calibration, and Imaging. The total time was calculated for 10 samples for each method. (**d**) Example image of a cleared T-47D spheroid labelled with DRAQ5 for nuclei (purple) and Phalloidin 488 for actin (green) and optically cleared with Sucrose protocol. A total of 5184 nuclei were segmented using *BIAS*. Scale bar represents 100 µm. (**e**) By co-culturing T-47D breast cancer cells with MRC-5 fibroblasts and EA.hy926 endothelial cells, a hydrogel-based tumour model was created to track the tumour microenvironment in real time. CellTracker dyes (orange, deep red, and green) are used to detect cells. The tumour, endothelial, and fibroblast cells are marked by red, green, and white labels, respectively. Scale bar represents 50 µm. (**f**) A ground truth dataset of HeLa Kyoto - MRC-5 co-cultures was created using the HCS plate. The dataset is available at https://doi.org/10.6084/m9.figshare.c.7020531.v1. Scale bar represents 50 µm. The Kolmogorov-Smirnov test was utilized for statistical analysis. *p ≤ 0.05; **p ≤ 0.01; ***p ≤ 0.001.

**Supplementary Fig. 3.**
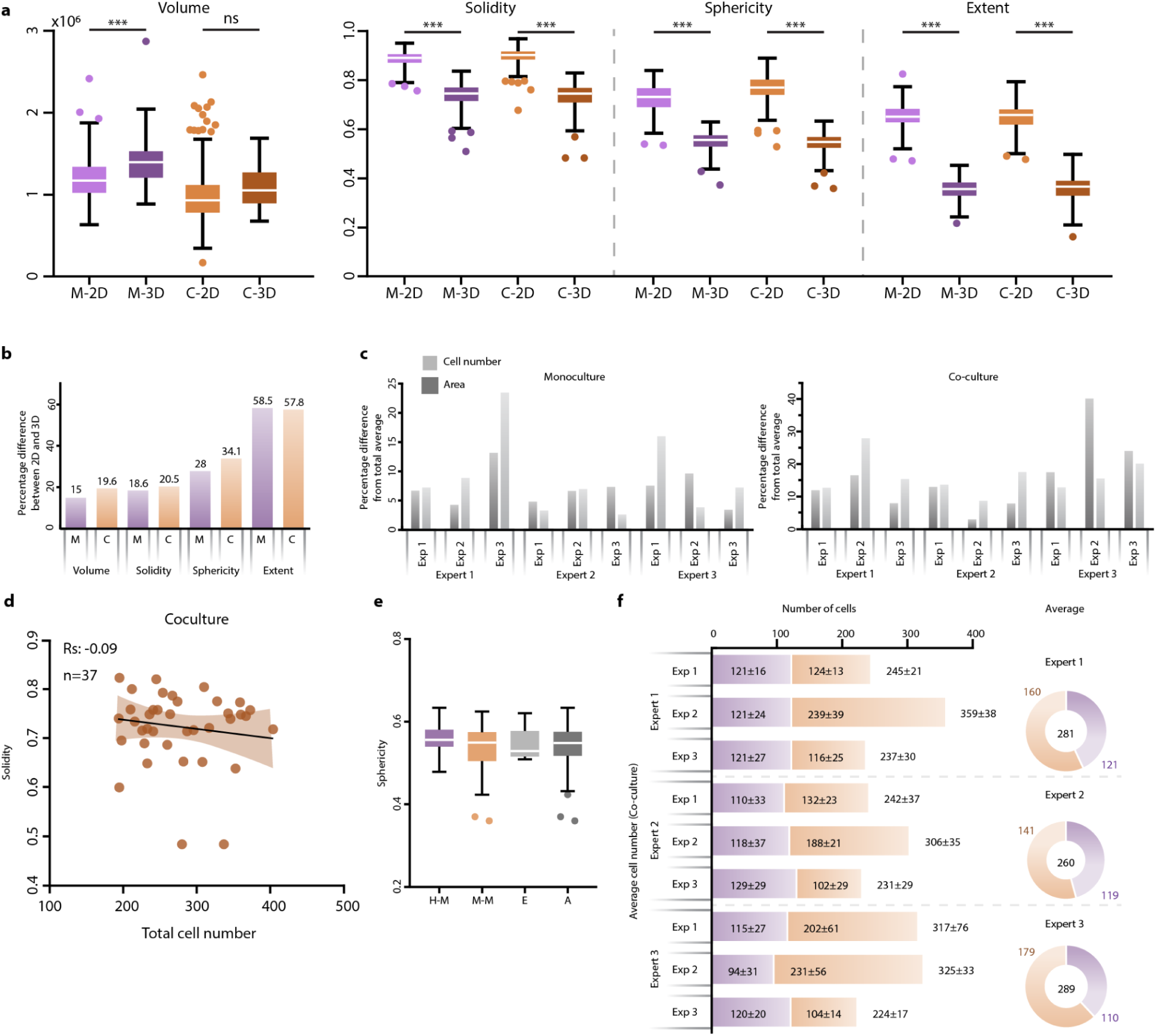
Boxplot (**a**) Percentage difference of the total average values for each experiment displayed for *Area* (dark) and total cell number (light). (**b**) Bar plot visualisation of the percentage difference from the average values between the 2D and 3D features. Purple colour represents the monoculture dataset, orange colour represents the co-culture dataset. (**c**) Barplot visualisation of the percentage difference from the total average of *Area* (dark grey) and *Cell number* (light grey) features. (**d**) Using the co-culture dataset, the Spearman’s correlation was measured between the total cell number and *Solidity*. (**e**) The 114 spheroids were divided into 3 groups based on the cell type majority (i.e. HeLa Kyoto majority, Equal, MRC-5 majority). *Sphericity* features collected from the spheroids were displayed based on the 3 groups and dark grey colour represents the average value of the whole dataset. (**f**) The average cell number of the 114 co-culture spheroids were displayed for each experiment (HeLa Kyoto - purple and MRC-5 - orange). The average value of cell number plated by each expert were visualised as a doughnut chart. The size of the doughnut charts and the numbers in their centre represent the average cell numbers for each group. A non-parametric Kruskal-Wallis test was conducted, followed by Dunn’s multiple comparison test for the statistical analysis. Kolmogorov-Smirnov test was utilized when only two groups were compared. *p ≤ 0.05; **p ≤ 0.01; ***p ≤ 0.001.

**Supplementary Fig. 4.**
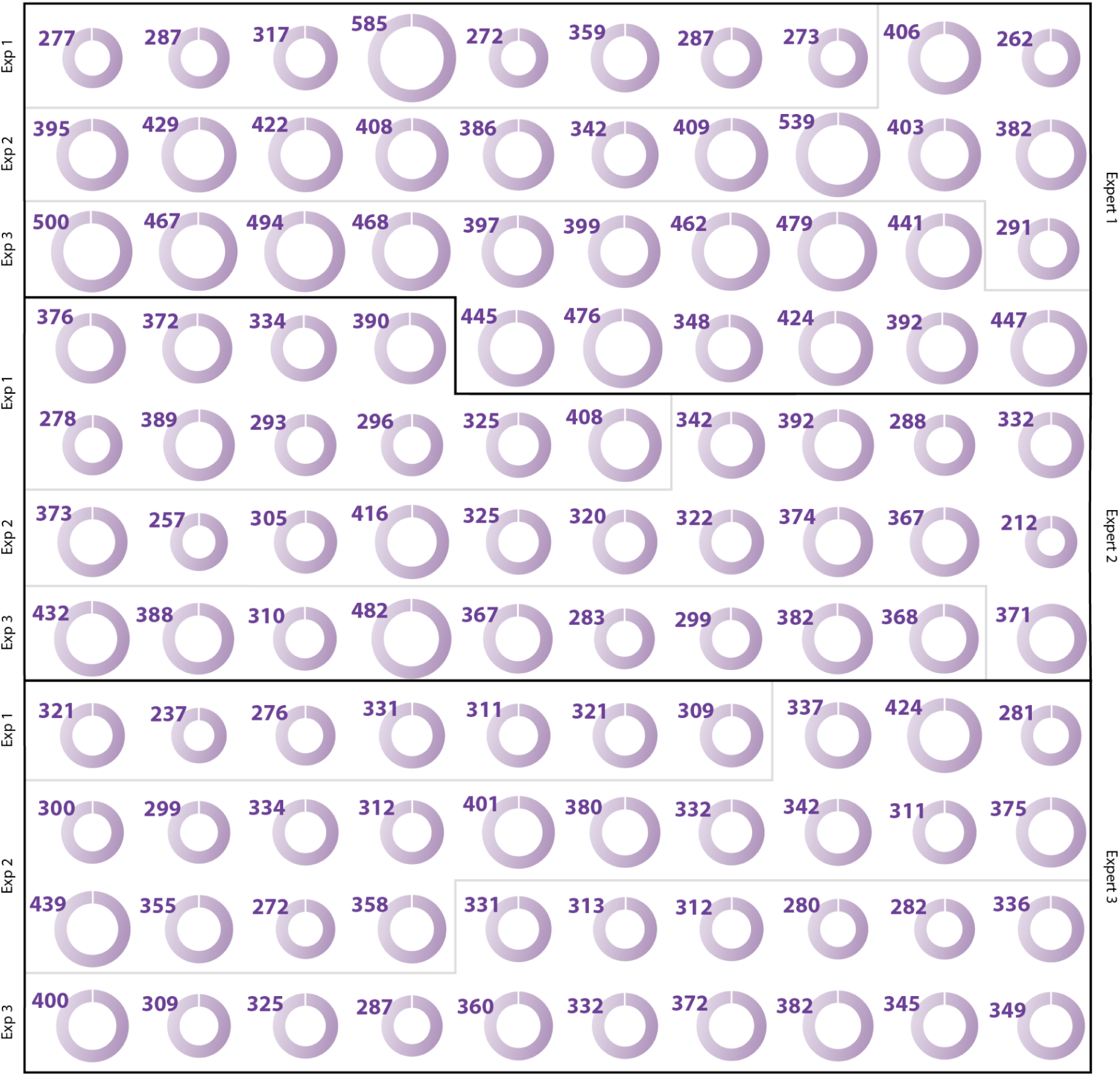
**(a)** Individual monoculture spheroids are displayed and separated by experts and experiments. The size of the doughnut chart refers to the total number of HeLa Kyoto cells (reported in purple); n=110.

**Supplementary Fig. 5.**
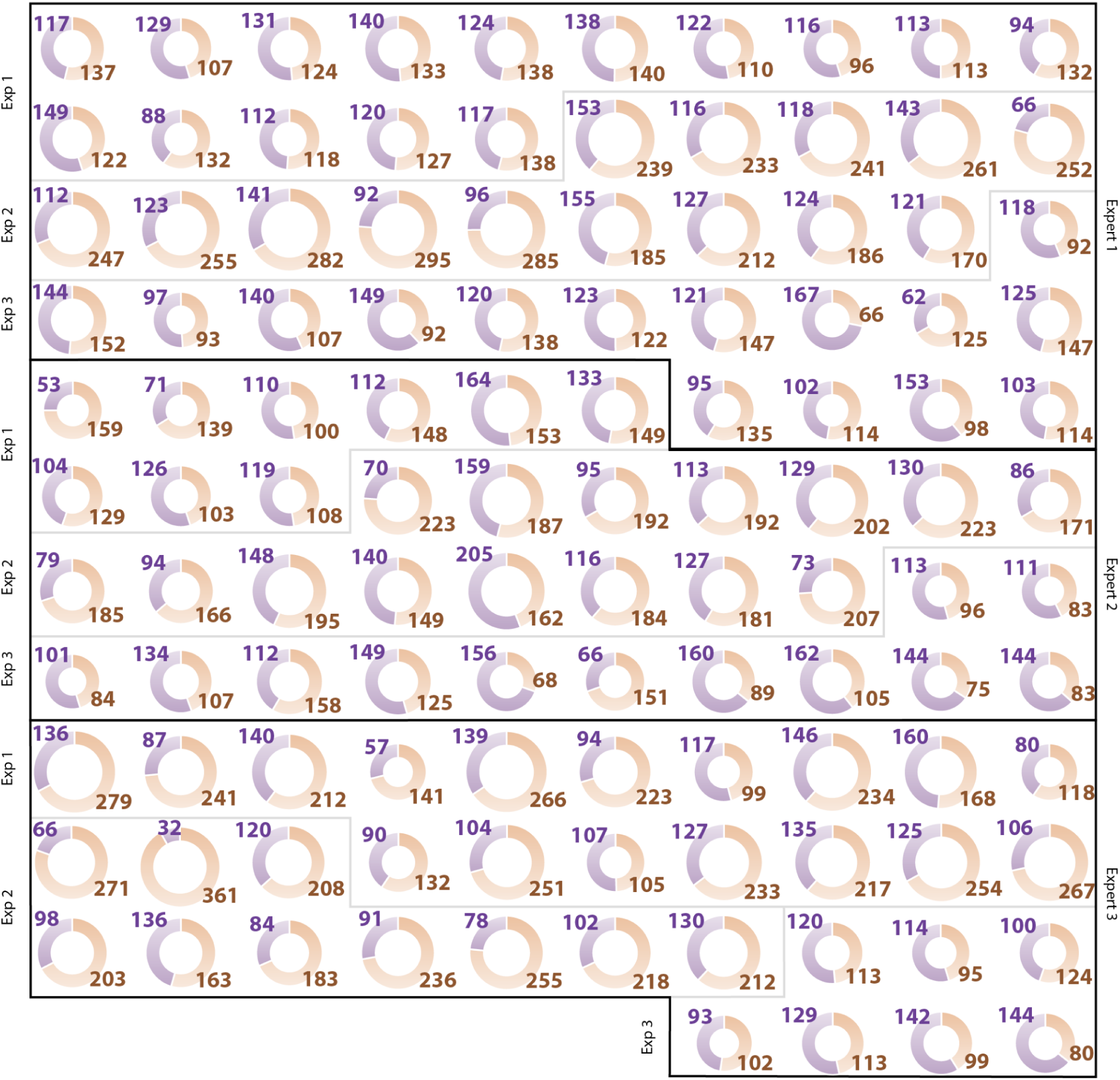
(**a**) Individual co-culture spheroids are displayed and separated by experts and experiments. HeLa Kyoto and MRC-5 cells are displayed with orange and purple colours. The size of the doughnut chart refers to the total number of cells (also reported in coloured numbers); n=114.

**Supplementary Fig. 6.**
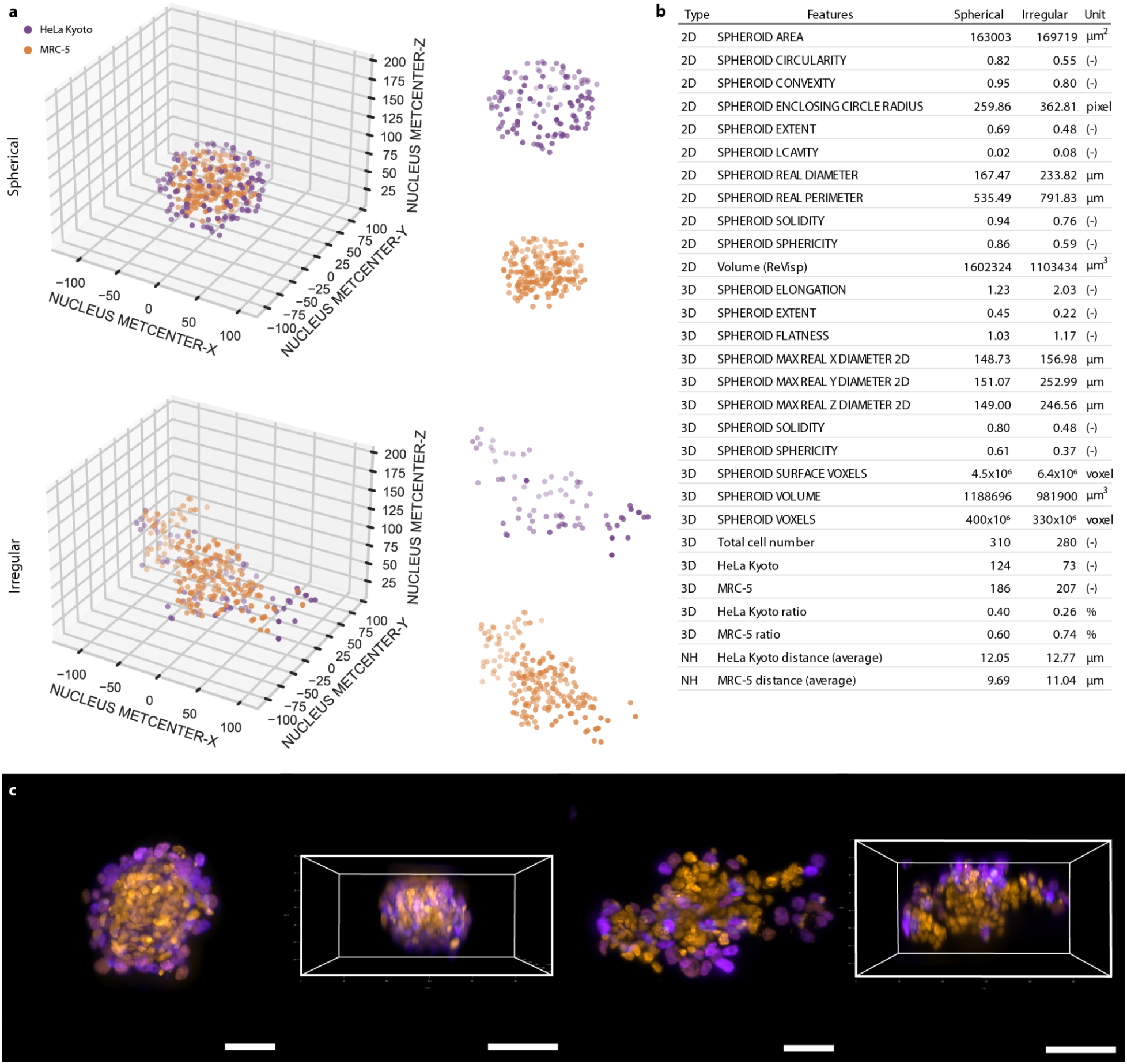
(**a**) 3D visualisation of the Spherical and Irregular spheroids where each circle represents a cell displayed based on their position within the spheroid and class (HeLa Kyoto - purple and MRC-5 - orange). Each class is also visualised separately. (**b**) Collected 2D, 3D, and Neighbourhood (NH) features for each spheroid are displayed. (**c**) 3D representation of the Spherical and Irregular co-culture spheroids. Both top and side views show the nuclei of the HeLa Kyoto (purple) and MRC-5 (orange) cells. For visualization, a square was placed around the spheroids indicating the side view. The scale bar represents 50 µm for the top and 100 µm for the side view. Leica’s LAS X microscope software was used to display the images.

## Supplementary Note 1

### HCS PLATE COMPONENTS

#### FEP foil

Fluorinated Ethylene Propylene (FEP) foils are transparent, with a refractive index of 1.341-1.347. FEP is chemically inert and resistant to most organic solvents, acids, and bases. Additionally, the material satisfies FDA (21CFR.177.1550) and EU (2002/72/EC) requirements (Hötte, K., *et al.*, Scientific Reports, 2019). FEP-films (Holscot Europe, Netherlands) with a 50 µm thickness were cut into 15 x 15 cm pieces and cleaned with 70% alcohol. Each foil was placed and clamped into the frame of a JT-18 vacuum-forming machine (Yuyao Jintai Machine Company, China). To get small and uniform shapes, the heater needs to raise the temperature to a level near the glass transition temperature of the FEP-foil which is between 260-280°C. Once the heater reaches the desired temperature, the positive mould is quickly placed onto the vacuum-forming machine, and the foil is quickly pressed onto the mould while the vacuum suction is switched on. The FEP foil that has been extruded is then carefully taken out of the mould and cleaned with an isopropyl alcohol bath for 5 min to remove the excess resin components. A 3D-printed holder was designed to provide support for the foil while transferring samples.

#### Insert element

To fix the samples’ position inside the cuvettes, hydrogels or a 3D-printed insert element are suitable (**Supplementary Note 1 Fig.1a)**. While hydrogels are optimal for smaller samples, the insert element offers an alternative solution to secure the position within the foil. The insert element contains the same pyramid-like structure as the foil, however, there is no sphere at the top. Inserting the element inside the cuvettes secures all samples in the spheres without deforming or harming the cells. To elevate the samples from the bottom to the same z-positions, the FEP foil with hydrogel or insert element should be turned over and placed inside of the base element. To provide precise positioning and a locking mechanism of the cuvettes inside of the base, the insert contains 4 magnets that attach to the 4 other magnets on the base.

#### Base element

The base components of the plate were printed with the Prusa i3 MK3S+ 3D printer with PETG (3DP-PETG1.75-01-BK; Gembird, Shenzhen, China) in two different size versions (**Supplementary Note 1 Fig.1b)** with the corresponding grid elements (**Supplementary Note 1 Fig.1c** and **d)**. The bigger version is 85 / 127 / 15.4 mm (average size of a 96-well plate) which is suitable for screening 56 samples, while the smaller version 85 / 73.8 / 15.4 mm was designed only for 28 samples. To create a waterproof base for the samples, the 24 / 64 mm coverslips (Menzel Gläser, Germany) were secured with silicone grease (ThermoFisher, USA) to the bottom of the bases. Models were designed in Blender and STL files (stereolithography files) were exported to *PrusaSlicer v 2.6.x*. All models were 3D printed with the default Prusa PETG filament profile and with “0.4 Quality print settings” using a 0.4 nozzle.

#### Grid element

To secure the foil’s position in the plate we printed a secure element that allows to fix the position of the FEP foil within the base (**Supplementary Note 1 Fig.1c** and **d)**. This compartment was printed with the Prusa i3 MK3S+ printer.

#### Positive mould

A positive mould for the spikes was created using *Blender 3.0*. Each mould was designed to fit perfectly into the base of the plate (**Supplementary Note 1 Fig.2a)**. For printing, the 3D models were exported to a stereolithographic file format (.stl) and then imported into *PrusaSlicer v 2.6.x* (Prusa Research, Czech Republic). Each heat-resistant mould was printed with the Prusa SL1 printer, which is a stereolithography (SLA) resin-based 3D printer that forms objects layer by layer using a liquid resin that is ultraviolet (UV)-cured. For printing, the DruckWege Type D High Temp resin (TDH-VIO-500, Groningen, The Netherlands) was used. This printer provides high-resolution printing capabilities enabling the production of intricate and detailed objects with smooth surface finishes. The mechanical and thermal characteristics of heat-resistant resin enable it to endure the vacuum-forming process. The positive moulds are UV-cured and washed with isopropanol to remove excess resin from the surface. Then the moulds were examined under a stereomicroscope and cleaned by submerging them in an ultrasonic bath before being used.

### ASSEMBLY OF THE HCS PLATE

First, biological samples should be placed precisely into the cuvettes of the FEP foil. An additional 3D printed element which is not part of the HCS plate can be used to prevent the movement of the foil while transferring the samples (***Supplementary Note 1 Fig.2c***). Prefilling the cuvettes of the foil with liquid before sample transfer can help remove air bubbles. The sample transfer process can be done manually, using a single or multi-channel pipette (8 channels) or a pipetting robot. After pipetting, a visual check is necessary because all the samples should sink to the bottom of the cuvette and positioned in the middle. If a sample is not in the middle of the cuvette, a gentle shake of the foil or pipetting the samples to the bottom may resolve this issue. If all the samples reach the bottom of the cuvettes of the FEP foil and there are no air bubbles next to the samples, then the position of the samples should be secured. Removing excess mounting media from the cuvettes allows the user to pipette a low melting point agarose and thus fix the position of the 3D sample inside the foil. The alternative solution is to gently push the insert element into the FEP foil until each cuvette is secured. By utilising either method, samples should be ready to be placed into the base element by turning over the whole FEP foil containing all the samples. At this point, all the samples should be at the top of the cuvettes covered by the transparent foil. Next, the FEP foil with the samples should be placed into the base element, where each cuvette is positioned within the coverslip. After checking the alignment of the cuvettes, the user must carefully fill the plate with the detection solution by removing all the air bubbles under the FEP foil. Before filling up completely, the grid element should be placed on top of the FEP foil to stabilise and prevent additional movements of the samples. Finally, before inserting the base into the microscope, the plate with the detection solution must be carefully filled to proceed with the calibration process (**Supplementary Note 1 Fig.2b**).

**Supplementary Note 1 Fig. 1.**
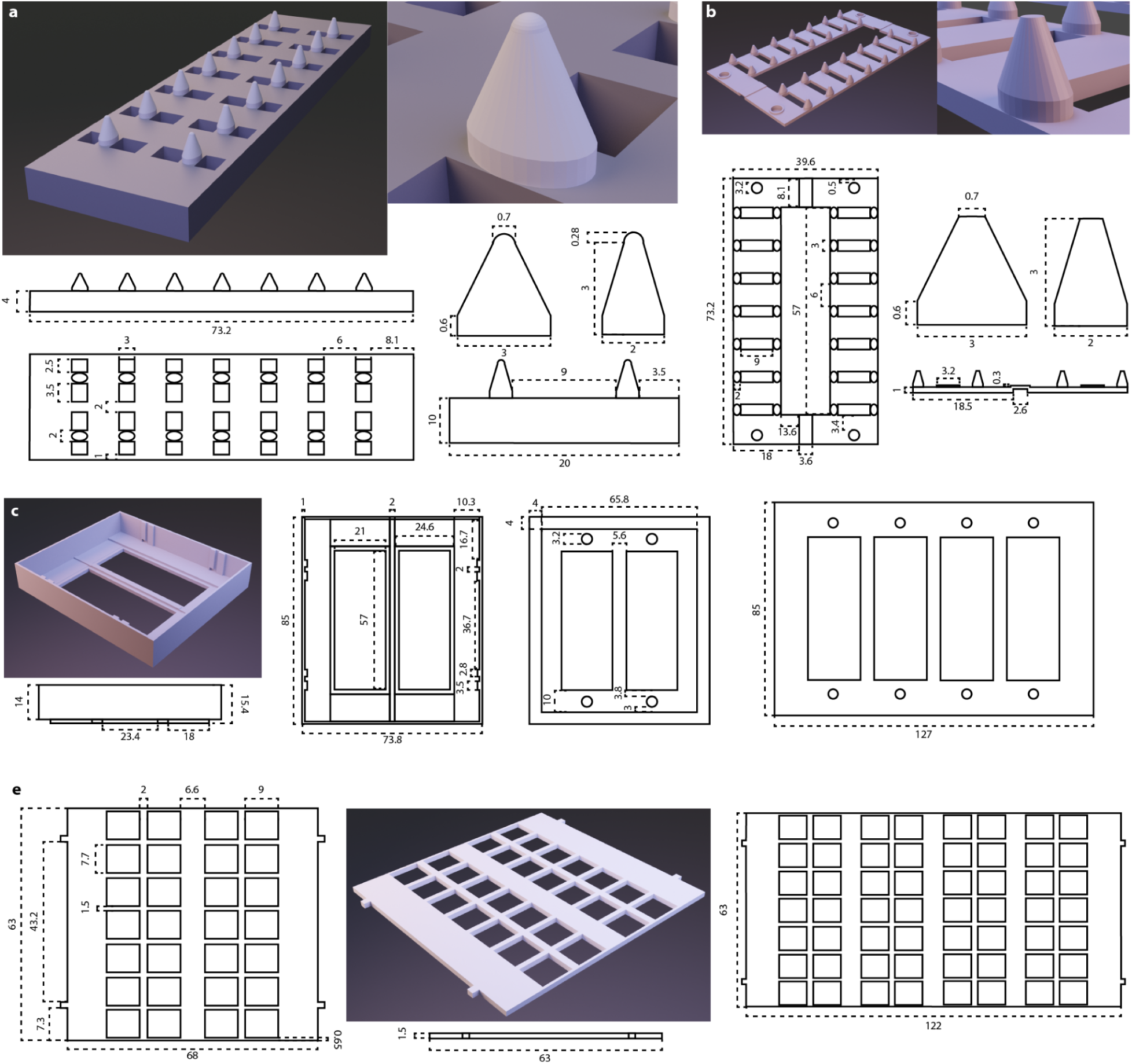
HCS Plate 3D printed elements (first part). (**a**) Positive mould element used for vacuum-forming, printed with the Prusa SL1 SLA resin-based 3D printer. (**b**) Prusa SL1 stereolithography (SLA) resin-based 3D printer was used to 3D print the insert element that secures the position of the samples within the vacuum-formed FEP foil. (**c**) The smaller and bigger version of the base element is designed for 28 or 56 samples within one plate. (**d**) Grid element that can be placed into the base element to fix the position of the FEP foil within the plate. The modified version of the grid element is shown that can be placed into the bigger base element. *Blender 3.0* software was used to create each rendered image. The dimensions of the figures are all displayed in mm. Every element was printed using a Prusa i3 MK3S+ printer except in cases where it is stated otherwise.

**Supplementary Note 1 Fig. 2.**
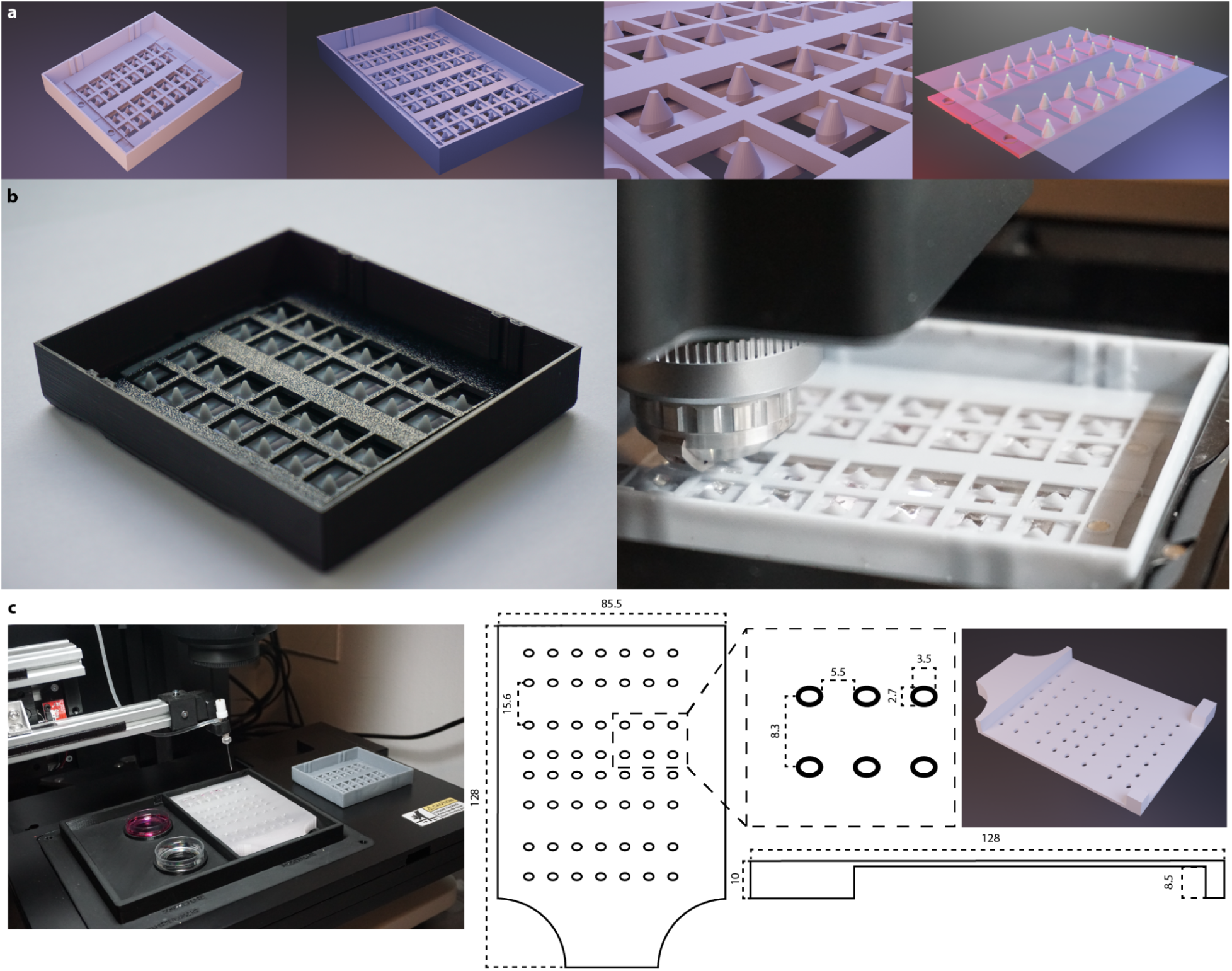
HCS Plate 3D printed elements (second part). (**a**) 3D rendered images representing the fully assembled plate for the smaller (28 samples/plate) and the bigger version (56 samples/plate). (**b**) 3D printed versions of the HCS plate. (**c**) An additional 3D printed element was made to prevent the moving of the vacuum-formed FEP foil while samples are transferred into the foil. This element is not part of the HCS plate and it is not needed for the assembly. It is compatible with *SpheroidPicker* and facilitates automated sample preparation. *Blender 3.0* software was used to create each rendered image. The dimensions of the figures are all displayed in millimetres. Every element was printed using a Prusa i3 MK3S+ printer.

## Supplementary Note 2

### 3D nuclei segmentation

For 3D nuclei segmentation, we employed the *StarDist-3D* method, a deep learning-based instance segmentation model designed for high precision in images with low signal-to-noise ratios and densely packed nuclei, such as spheroid images (**Supplementary Note 2 Fig. 1a**). *StarDist* predicts cell shapes using a star-convex polyhedron representation, directly estimating the shape parameters for each pixel and applying non-maximum suppression to ensure unique object predictions.

To train the model, we prepared an annotated dataset of 21 3D images of fluorescently labeled nuclei, resulting in 7238 manually and semi-automatically annotated nucleus instances. Image data for the Embryo and Annotator datasets (https://doi.org/10.6084/m9.figshare.c.7020531.v1) and for the optically cleared spheroids (https://doi.org/10.6084/m9.figshare.c.5051999.v1) are freely available (**Supplementary Note 2 Fig. 1b**). We resized the annotated images to get a median nuclei size of 68 × 68 × 5 voxels. We used a patch size of 256 × 256 × 15 voxels to guarantee that the network’s field of view could encapsulate at least one nucleus. Data augmentation techniques, including random rotations, flips, and intensity adjustments, were applied during training to enhance robustness. The dataset was split into 15 training and 6 validation images. Training was conducted for 200 epochs on an NVIDIA Titan Xp GPU with 12 GB of graphical memory, requiring approximately 3 days.

Performance evaluation followed the metric shown in **Supplementary Note 4 Eq. 4,6**. With an IoU threshold set to 0.7, our trained model achieved a precision of 0.6453. Prediction of a complete 3D image stack at their native resolution (2048 x 2048 x 60) required approximately 7.5 mins.

### Whole spheroid segmentation

To segment entire spheroids, we trained a 2D *U-Net* model using the fastai Python package. For training data, we annotated 10 3D images of spheroids and utilized their individual z-slices as 2D images. This approach allowed us to leverage 2D convolutional networks for efficient training while capturing the essential features of the 3D structures. The dataset was randomly split into 80% for training and 20% for validation, and the model was trained for 100 epochs.

Once trained, the model was applied to predict spheroid regions in new images. As a post-processing step, we performed a hole-filling operation to ensure continuity of segmented regions and removed small, disconnected objects that were not part of the spheroids. To evaluate performance, we used the metrics outlined in **Supplementary Note 2 (Eq. 4,6)**. Setting the IoU threshold to 0.7, the segmentation results demonstrated a precision of 0.7538. Segmentation on new images was performed on an NVIDIA GeForce RTX 3060 GPU with 6 GB of graphical memory, achieving an average processing time of approximately 150–200 milliseconds per 2D image. This pipeline enabled robust and accurate segmentation of whole spheroids for further analysis.

To provide a more accurate comparison for the experiments, each mask was checked and manually corrected by an expert. Measured features were compared before and after the mask correction, however, no significant differences were measured (**Supplementary Note 2 Fig. 1c**).

**Supplementary Note 2 Fig.1.**
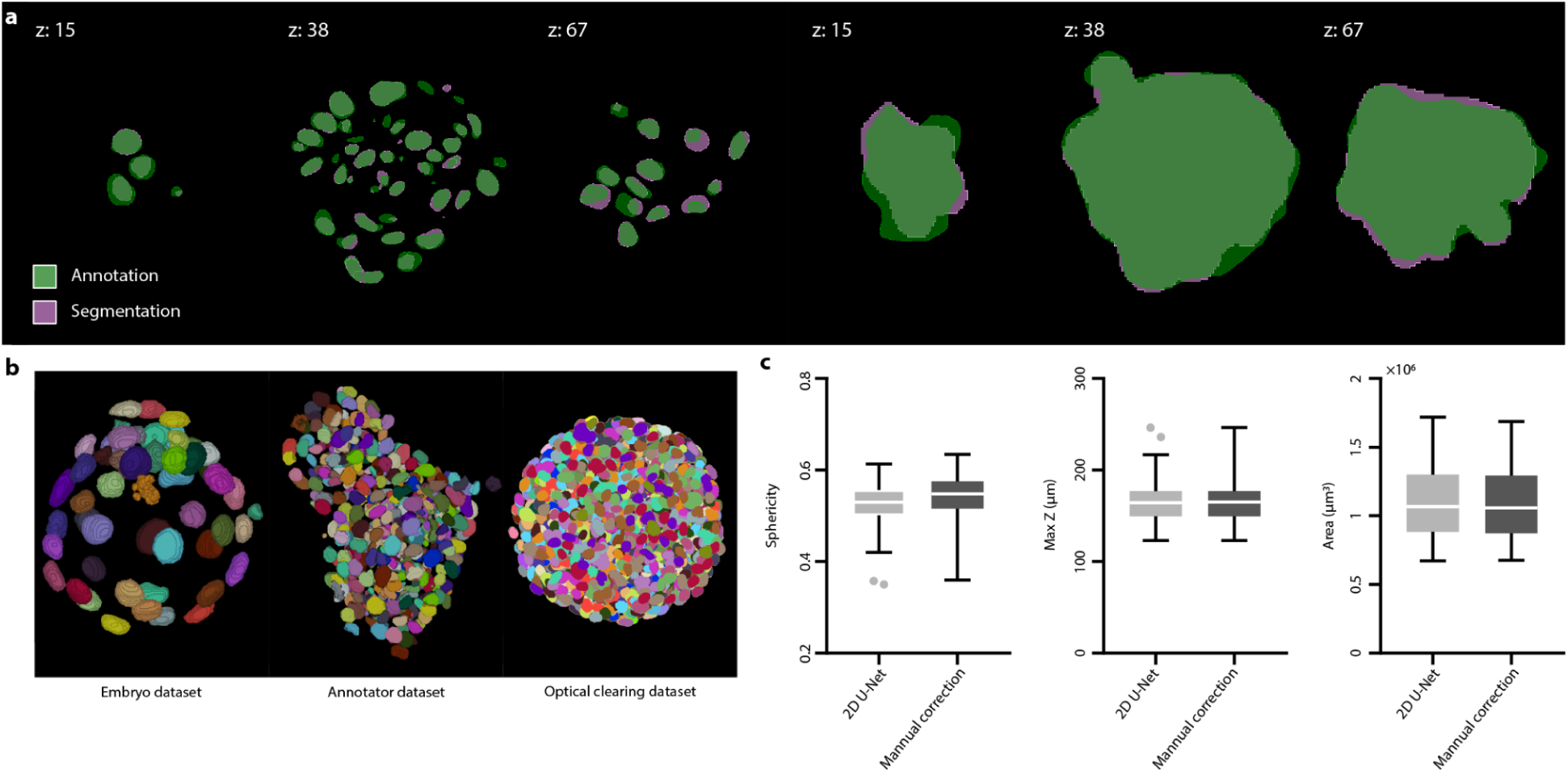
Segmentation. (**a**) Comparison of the accuracy of the *StarDist* and 2D *U-Net* models used on an annotated co-culture spheroid. Images with their corresponding masks were selected from the top (z: 15), middle (z: 38), and bottom (z: 67) regions of the spheroid. The first 3 images on the left show the nuclei segmentation result acquired by the *StarDist* model, while the following 3 images belong to the whole segmentation that was achieved by using 2D *U-Net*. Result of the segmentation is represented with purple and the manual annotation is represented with green. (**b**) Representative images of the Embryo, Annotator, and Optical clearing datasets used for the training of the *StarDist* model. (**c**) Boxplot visualisation of image-based features (*Sphericity*, *Maximum Z distance*, and *Area*) showing the result of 2D *U-Net* whole spheroid segmentation compared to manual correction. Data was obtained by evaluating 114 co-culture spheroids.

## Supplementary Note 3

All 2D and 3D raw images were saved as an *.xlef file format using the *LAS X* software (Leica, Germany), then images were directly imported into the *BIAS* software (Single-Cell Technologies Ltd., Szeged, Hungary). Due to the nature of the data, different workflows were created for the 2D and 3D datasets. Therefore, these were handled separately during the analysis.

### 2D workflow

The *AnaSP* software was used to manually annotate brightfield images in order to obtain the *ReViSP* based volume features (*e.g. Volume 2D*) and binary masks. The original images with their corresponding masks were imported into *BIAS* software. The import process is the only manual task that is required in the workflow, the rest of the pipeline can be executed automatically. To import and visualise the segmentation results, the *Segmentation* module was used. Next step is the *List creator*, an intermediate module that links results of different segmentations into a list of meta-objects that serve as an input for the future processing steps in the pipeline such as the *Feature Extraction* module that extracts various features, like intensity, shape, size, etc. of these meta-objects (spheroid in this case). Using the *Exporters* module, all features were exported as a *.csv file.

### 3D workflow

For the 3D dataset, a *StarDist* and a 2D U-Net models were used to segment nuclei (using only the blue channel - DAPI) and the whole spheroid (using only the gray channel - actin) in 3D using the *Execute module* (**Supplementary Note 3.**). This module allows the execution of third-party applications or scripts as a part of a *BIAS* pipeline. Generated masks were visualised in the *3D segmentation* module. *Mask operator/exclude* function was used to discard all objects outside of the spheroid’s volume and keep objects on the border and within the spheroid. In case of the nuclei mask, the *Mask operator/dilate* function with a 4 µm dilatation was utilised to increase the volume and to collect cell information. *Object linker* module has a similar functionality as the *List Creator* and it was used to create the meta-objects with the corresponding spheroid, cell, and nuclei information and the generated output was used to collect features with the *Feature extraction* 3D. By using a supervised machine learning algorithm from the *Machine learning classification* module, cells of the co-culture spheroids were classified either as a HeLa Kyoto or an MRC-5. To distinguish the cells’ type, the intensity channels 2 and 3 (green - EGFP-alpha-tubulin; red - H2B-mCherry) were used to create a training dataset with 750 classified objects in each category. Multi-Layer Perceptron (MLP) classifier was used to classify all the cells with a combined average accuracy of 98.59% according to a K-Fold cross validation (K=10) (where classes reached 98.40% for the HeLa Kyoto and 98.78% for the MRC-5 classes). Using the *Statistics* module, segmented objects with a particularly small size and missing features were removed from further analysis (NUCLEUS CONTOUR STAT MEAN > ‘20’ and CELL CONTOUR STAT MEAN > ‘30’). To measure the closest distance between cells, *Neighborhood features/K-neighbors* ‘1’ was used for both classes separately. Finally, all features were exported as a *.csv file using the *Exporters* module.

**Supplementary Note 3 Fig.1.**
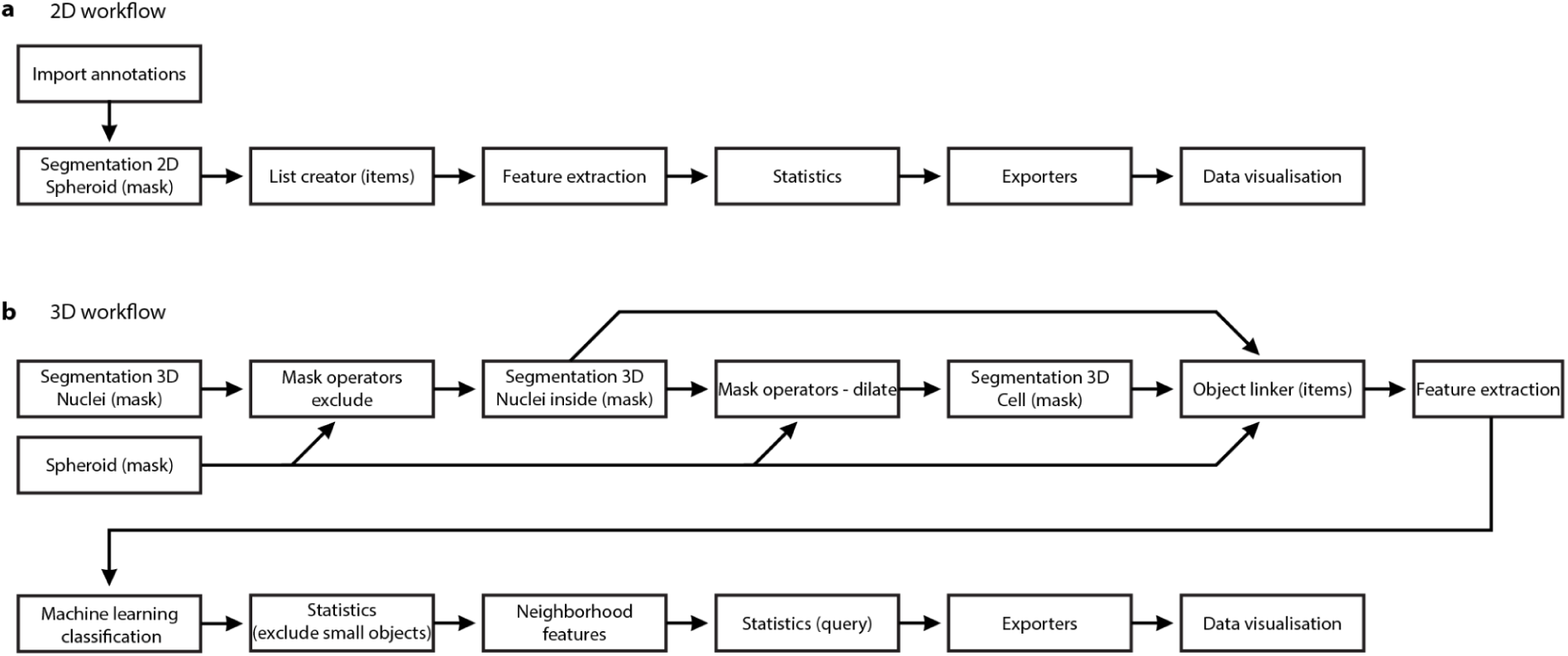
*BIAS* workflows for data analysis. (**a**) 2D workflow to evaluate brightfield images of spheroids. (**b**) 3D workflow of spheroid dataset imaged by LSFM.

## Supplementary Note 4

### SpheroidPicker

According to our findings discussed in this article, we improved and updated *SpheroidPicker* that was published previously^18^. As a result, new features were added to the pre-selection of spheroids, the GUI was updated to enable user-friendly selection, the AI-based image detection model was enhanced, and the HCS plate was added as a selectable option.

### Software

For the 2D pre-selection, the original method provided only limited options for feature extraction of the -oids (*Area*, *Perimeter*, and *Circularity)*. To set criteria for sample selection, the new version contains features that may better approximate the 3D properties of candidate objects (**Fig. 3.g and h**). These features are the *Maximum diameter*, *Sphericity*, and *Solidity* (**Supplementary Note 4 Eq: 1-3**) and are defined as:

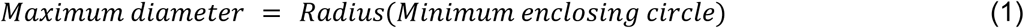

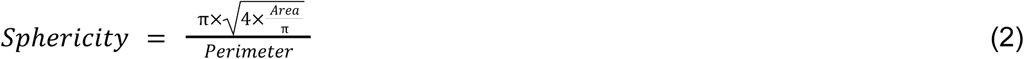

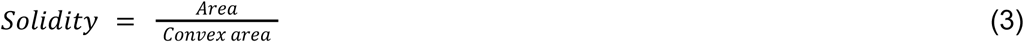

Where *Area*, *Perimeter*, *Convex area*, and *Minimum enclosing circle* were calculated utilizing OpenCV libraries. The GUI now allows the selection of spheroids based on the updated features or their arbitrary combinations and offers the export of calculated features.

### Segmentation model improvements, benchmark and dataset extension

In order to show the robustness of the models, we have created a new test set that is completely unseen by the model. 134 Brightfield images were acquired using the stereomicroscope’s (Leica S9i, Germany) integrated 10 megapixel camera. Ground truth annotations were created by using the AnnotatorJ*/ImageJ* plugin (https://doi.org/10.1091/mbc.E20-02-0156).

To improve the performance and generalizability of the *Mask R-CNN* model for spheroid segmentation, photometric and geometric augmentations were used. The following transformations were applied randomly on the training image set: translation, scaling, cropping, rotation, flip, brightness, contrast, sharpness, adding noise, inversion, and color channel shifting. The transformed images were added to the training set. The training was performed with the same parameters as reported previously^18^. The original and the improved performance of the models were compared using precision and recall metrics on different Intersection over Union (IoU) thresholds (**Supplementary Note 4 Eq. 4-6**).

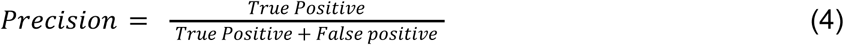

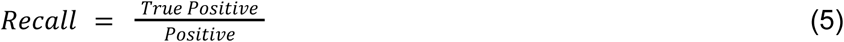

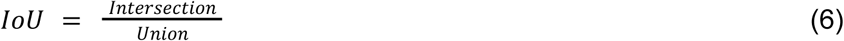

First, we calculated the True Positive (TP) and False Positive (FP) values for IoU thresholds ranging from 0.5 to 0.95, with increments of 0.05. Using these values, we computed the precision and recall scores at each threshold (**Supplementary Note 4 Fig 1**).

**Supplementary Note 4 Fig. 1.**
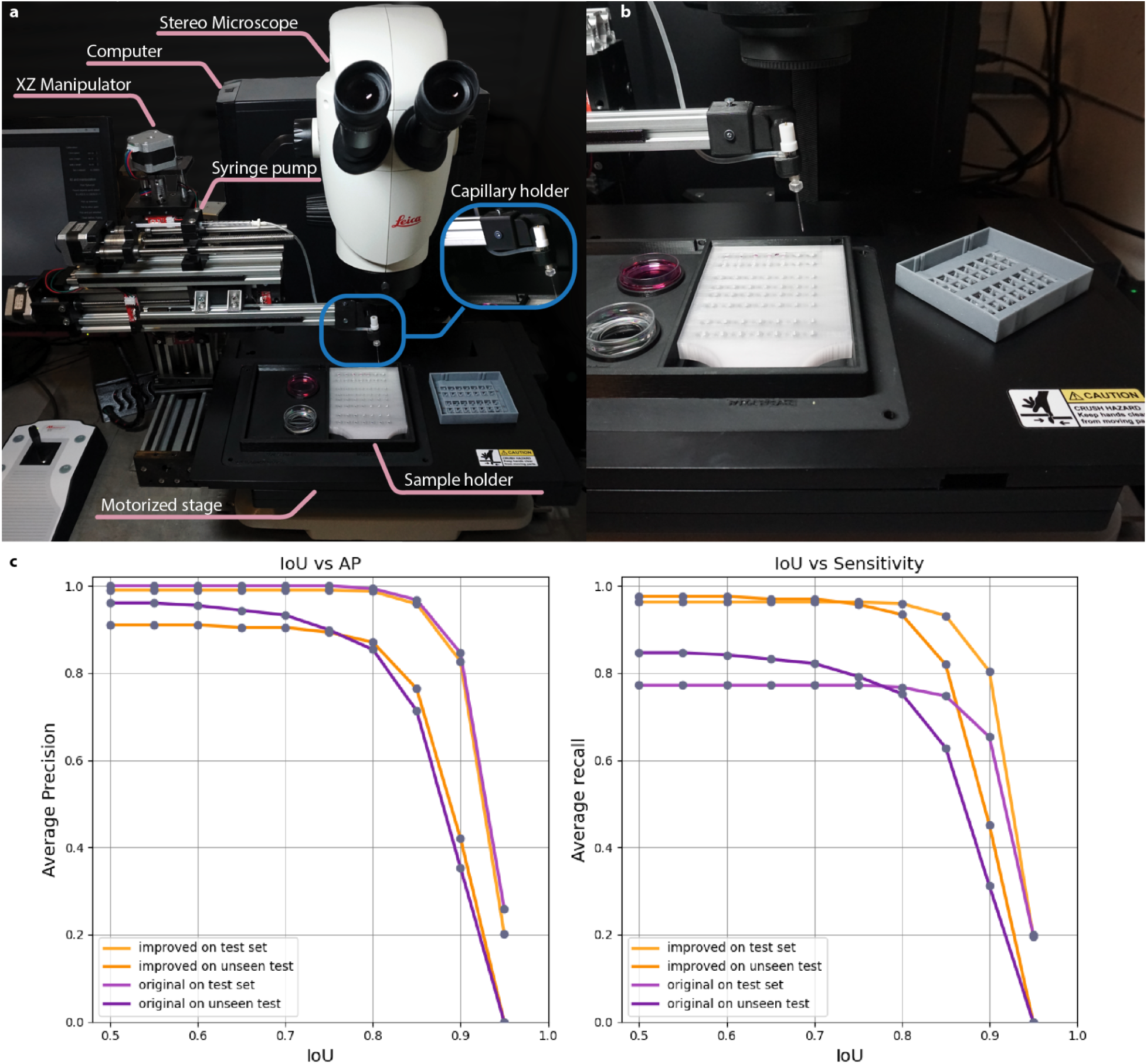
Improved version of the *SpheroidPicker*. (**a**) Concept of the SpheroidPicker. The image displays all the components required to transfer samples from a source plate into a target plate. (**b**) Representative image of transferring spheroids into the FEP-foil. (**c**) Comparison of the segmentation methods, average precision and recall at different intersection over union (IoU) thresholds. The performance of the previously used models are displayed in light and dark purple (test set and unseen test), while the new versions are labeled with light and dark orange (test set and unseen test).

Precision scores are nearly equal between the new and original models on both the test and unseen test sets. However, the recall curve shows that the augmented model achieves higher scores, indicating that it can detect more objects compared to the original model.

We made the source code of SpheroidPicker opensource on github https://github.com/grexai/SpheroidPicker. The model files are also available, to 3D print at (https://doi.org/10.5281/zenodo.14679243). The original and the improved segmentation models are available at zenodoo (https://doi.org/10.5281/zenodo.14675683), and finally the annotated dataset (https://doi.org/10.5281/zenodo.14679303).

